# An iris-like pore for protein import into peroxisomes

**DOI:** 10.64898/2026.10.23.761396

**Authors:** Dingwei He, Yuan Gao, Tom A. Rapoport

## Abstract

Peroxisomes are ubiquitous, membrane-bound organelles that house essential metabolic enzymes imported from the cytosol in a folded and sometimes oligomeric state. Mobile receptors, such as PEX5, ferry cargo through a tyrosine/glycine (YG)-rich selective phase formed by PEX13 within the membrane, but how folded proteins of different sizes cross the membrane remains unknown. Here we report structures of the channel that mediates cargo import. Twelve PEX13 molecules assemble into a pore formed by a ring of tilted amphipathic helices in alternating orientations. Hydrophobic faces contact the lipid bilayer, whereas hydrophilic faces line the pore containing the YG phase. Pairs of oppositely oriented helices can slide relative to neighboring pairs and thereby alter the helix tilt angle, enabling iris-like dilation of the pore from ∼52 Å in the resting state to ∼70 Å, sufficient to accommodate the PEX5 receptor carrying small cargo. Translocation of the largest cargo requires further widening to ∼100 Å, a state supported by AlphaFold modeling and disulfide crosslinking. Our findings suggest that the channel provides an adaptive conduit for folded proteins of varying sizes, a mechanism distinct from other known modes of protein translocation.

## Main text

Peroxisomes are ubiquitous organelles that consist of a single membrane enclosing a lumen (matrix) packed with proteins involved in various metabolic pathways, including fatty acid oxidation and detoxification of reactive oxygen species. Peroxisomes are vital for human health; peroxisomal disorders lead to debilitating and often fatal diseases, notably the Zellweger spectrum disorders (for review, see ^1–4^).

Peroxisomes import their matrix proteins post-translationally from the cytosol (for review, see ^1,2,5–9^). Remarkably, these proteins are translocated across the membrane in a folded or even oligomeric state, unlike proteins destined for the ER, mitochondria, or chloroplasts, which must be unfolded for import. How folded proteins of different sizes traverse the peroxisomal membrane remains unknown.

Peroxisomal protein import is mediated by soluble receptors that recognize cargo in the cytosol and deliver it to the matrix. Most imported proteins carry a C-terminal peroxisome-targeting signal (PTS1), typically the sequence SKL or a close variant ^10^, recognized by the tetratricopeptide-repeat (TPR) domain of the receptor PEX5 ^11^. The cargo-bound receptor is recruited to the peroxisomal membrane by the “docking complex”, which contains the conserved membrane proteins PEX13 and PEX14 ^12^. The receptor then ferries bound cargo through the peroxisomal membrane all the way into the matrix ^13^. Translocation occurs through a nuclear pore-like conduit formed by PEX13 ^14,15^. PEX13 contains a conserved tyrosine/glycine-rich (YG) domain that resembles the phenylalanine/glycine (FG) repeats of nucleoporins ^16–18^. The YG domains of multiple PEX13 molecules form a meshwork within the membrane, generating a selective phase that excludes inert molecules above a size threshold while allowing receptors to partition into it through conserved aromatic motifs in their unstructured N-terminal regions ^14^. After cargo delivery, the receptor is recycled to the cytosol by a retro-translocon comprising a membrane-embedded ubiquitin ligase complex, followed by extraction by a hexameric ATPase ^19–22^. A subset of matrix proteins carries an N-terminal PTS2 signal that binds to both a receptor and the adaptor PEX7 ^23,24^. This trimeric complex traverses the YG phase to deliver cargo to the matrix; the receptor is then recycled by the retro-translocon, while PEX7 returns independently to the cytosol through the YG conduit ^25^ and is extracted by PEX39 ^25,26^.

The central unresolved question concerns the architecture of the import conduit itself. Recent work suggested that PEX13 forms the translocon for peroxisomal import ^14^. PEX13 has only two conserved features: the unstructured YG domain and a long amphipathic helix (AH) (**Fig. 1a**). The YG domain is preceded by an additional disordered region that is only required for PTS2 import ^25–27^. In many organisms, PEX13 also has a predicted C-terminal transmembrane (TM) segment and an SH3 domain (**Fig. 1a**), but these are absent in plants and algae ^8^. Protease-protection experiments indicate that PEX13 adopts dual topology in the membrane: half the molecules have their C terminus facing the cytosol and the other half facing the matrix ^14^. Although dual topology is unusual among membrane proteins, it allows the YG domains of both PEX13 populations to meet within the plane of the membrane and form a meshwork. How this YG phase is suspended within the membrane and insulated from the surrounding lipid remains unknown.

**Fig. 1.**
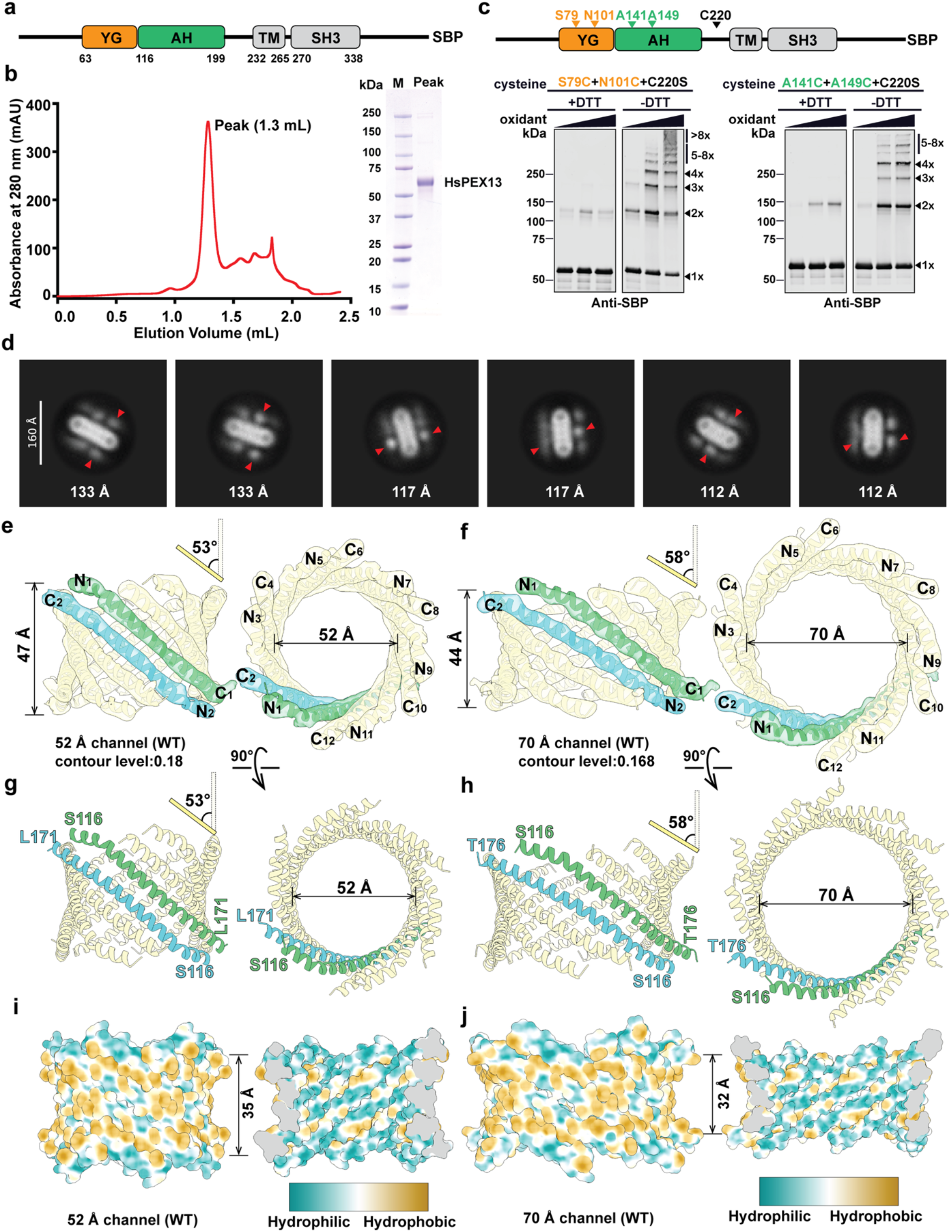
Cryo-EM structures of human PEX13. **a**, Domain organization of human PEX13. **b**, SBP-tagged PEX13 was expressed in *P. pastoris*. The protein was solubilized in digitonin and purified on streptavidin beads followed by SEC (left panel). The peak fraction was analyzed by SDS-PAGE and Coomassie blue staining (right panel). **c**, PEX13 containing two cysteines at the indicated positions of the YG or AH domains and with the endogenous cysteine 220 mutated to serine was purified as in (B) and treated with Cu-phenanthroline (oxidant). The samples were analyzed by nonreducing (–DTT) or reducing (+DTT) SDS-PAGE and immunoblotting with SBP antibodies. The number of crosslinked PEX13 molecules in the bands is indicated on the right. **d**, Purified human PEX13 was subjected to cryo-EM single-particle analysis. Shown are sideviews of 2D class averages. Arrowheads indicate SH3 domains on both sides of the detergent micelle. Note the variable lengths of the particles. **e**, Semi-transparent density map of the ∼52 Å channel in two views, with a docked cartoon model based on AlphaFold3 (**Extended Data Fig. 4a**). Two of the 12 AHs are highlighted. The helix tilt angle is shown on top of the left panel. **f**, As in **e**, but for the ∼70 Å channel. Another particle class has small C-terminal extensions of the AHs (**Extended Data Fig. 3a**). **g**, Model for the AHs of the ∼ 52 Å channel. **h**, As in **g**, but for the ∼70 Å channel. **i**, Space-filling model of the ∼52 Å channel. The left panel shows the outside of the pore, and the right panel the inside in a cut-away view. Hydrophobic and hydrophilic residues are colored with the scale on the right. The size of the hydropobic belt is indicated. **j**, As in **i**, but for the ∼70 Å channel.

Here we report cryo-EM structures of the peroxisomal protein import channel. Twelve PEX13 molecules assemble into a pore ring in which amphipathic helices form the channel wall around the YG phase. Experimental structures of 52 Å and 70 Å pores, together with a predicted 100 Å pore supported by disulfide crosslinking, show that helix pairs slide relative to one another, allowing the channel to dilate like an iris and accommodate folded cargo of different sizes.

### Cryo-EM structures of the wild-type PEX13 channel

To obtain a structure of the PEX13 channel, we expressed the wild-type human protein with a streptavidin-binding peptide (SBP) tag at the C terminus in *Pichia pastoris* cells (**Fig. 1a**). A membrane fraction was isolated, the protein was solubilized in the detergent digitonin and purified by binding to streptavidin beads followed by size-exclusion chromatography (SEC) (**Fig. 1b**). The protein eluted as a sharp peak at high molecular weight. Analysis of the peak fraction by SDS-PAGE and Coomassie blue staining revealed a single band of expected size (**Fig. 1b**). PEX13 maintains its oligomeric state after detergent solubilization and purification, as shown by introducing two cysteines into either the YG or AH domain and adding an oxidant. In both cases, a ladder of disulfide-linked adducts was observed, which largely disappeared after reduction of the disulfides with dithiothreitol (DTT) (**Fig. 1c**).

The purified wild-type (WT) protein was used for cryo-EM single-particle analysis. 2D classification of the particles showed side views in which the detergent micelle is bordered by several dots (arrowheads in **Fig. 1d**; **Extended Data Fig. 1a-c**). These should correspond to the SH3 domains, as PEX13 does not possess any other globular domain outside the membrane. Thus, in agreement with the predicted dual topology in intact membranes ^14^, some PEX13 molecules have their SH3 domain on one side of the detergent micelle, while others have it on the opposite side. The side views also indicate that the dataset contains particles of different sizes (**Fig. 1d**).

Consistent with the heterogeneity of the particles, we obtained different density maps from the dataset with 6-fold symmetry imposed during data processing (**Extended Data Fig. 2; Extended Data Table 1**). All reconstructions resulted in refined density maps with an overall resolution of ∼6.2 Å (**Fig. 1e; f; Extended Data Fig. 3a**). The maps show a ring of twelve left-tilted helices surrounding an almost circular pore. In two maps, the pore diameter is ∼70 Å, and in the third ∼52 Å. The two density maps with the larger pore size are almost identical; the only difference is additional density at the end of the helices (**Extended Data Fig. 3a**). Because of the limited resolution, no helical turns or side chain density were discernible.

The tilt angle of the AHs (defined as deviation from the membrane normal) is smaller for the ∼52 Å pore than the ∼70 Å pores (53° versus 58°) (**Fig. 1e; f**). The channels also differ in their height (47 Å versus 44 Å). The SH3 domains were seen at low contour level (**Extended Data Fig. 3b**), but seem to be flexible. The TMs might be responsible for the weak density seen at the periphery of the pore (**Extended Data Fig. 3c**).

Molecular models for the ∼52 Å and ∼70 Å channels were derived with the help of AlphaFold ^28^. Using 12 copies of the AHs, AlphaFold3 predicts channels of different diameters with low confidence scores (ipTM 0.2–0.3). Including 12 copies of the TMs in the analysis, resulted in improvement of the predictions, although AlphaFold3 randomly converged on channels of either ∼52 Å or ∼70 Å diameter (**Extended Data Fig. 4a**). In both cases, the AHs were predicted with much higher confidence than the TMs and formed pores that fit well into the experimental density maps (**Fig. 1e-h**). Models for the AHs start at the N-terminus of the AH (residue S116) and extend to residues 171 and 176 in the ∼52 Å and ∼70 Å channels, respectively (**Fig. 1g; h**). Residues 177 to 199 of the AH are invisible in all maps. A kink in the density map is seen at opposite ends in neighboring helices, consistent with the AlphaFold models in which AHs alternate in their membrane topology (**Extended Data Fig. 4a**), and with the placement of the SH3 domains to both sides of the channels (**Extended Data Fig. 3b**). Two AHs of opposite orientation form a unit that is repeated six times in the certain tilt angle around the ring, explaining why imposition of 6-fold symmetry was required for structure determination. When viewed perpendicular to the membrane plane, the helices are curved, forming blades as in an iris diaphragm (**Fig. 1e; f**). In both channels, the AHs have their hydrophobic faces oriented towards the lipid and their hydrophilic faces pointing towards the interior of the channel (**Fig. 1i; j**), supporting the validity of the AlphaFold predictions. Furthermore, all helix interfaces leave only small holes in the channel walls.

As helix polarity was difficult to assign from the density maps, we used *S. cerevisiae* PEX13 to confirm that neighboring AHs in the PEX13 channel have opposite membrane orientations. Endogenous yeast PEX13 with a single cysteine in the AH was fused at its C terminus to a 3C protease-cleavage site, followed by an SBP tag to enhance the size shift after proteolysis. An N-terminal FLAG tag was added for immunodetection (**Fig. 2a**).

**Fig. 2.**
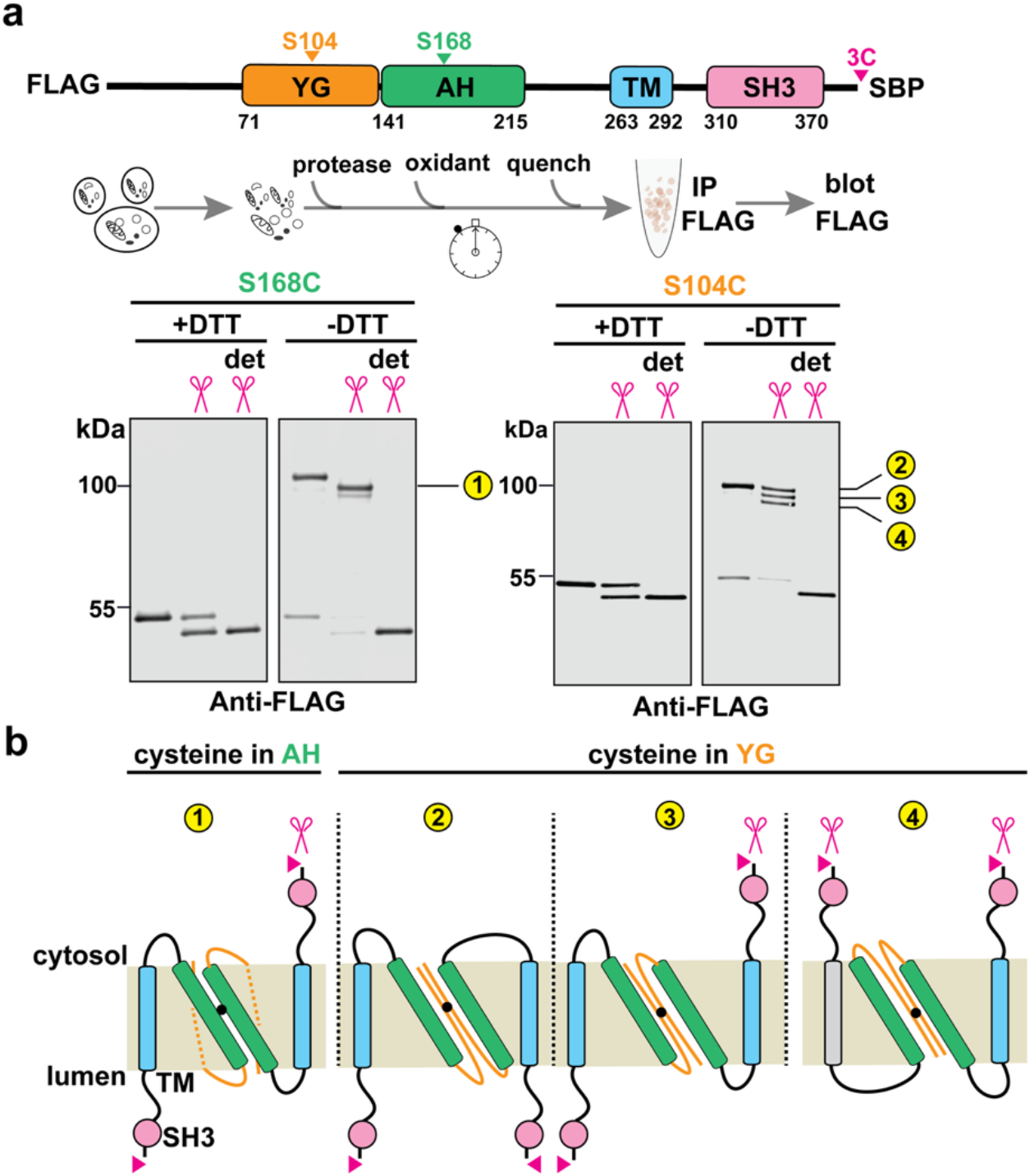
The AHs of *S. cerevisiae* PEX13 have alternating orientations. **a**, A cysteine was incorporated into the AH (green) or YG domains (orange) of FLAG-tagged yeast PEX13 containing a C-terminal 3C protease–cleavage site and SBP-tag, as indicated. The SBP tag serves to increase the size shift after proteolysis. Intact yeast membranes were treated first with the protease and then with the oxidant Cu-phenanthroline. The samples were quenched, and the proteins were immunoprecipitated (IP) and analyzed by nonreducing SDS-PAGE and immunoblotting with FLAG antibodies. Where indicated, dodecylmaltoside (det) was included during protease cleavage. The numbers refer to the topologies of the observed dimers (schemes in (**b**)). **b**, With the cysteine in the AH, only one crosslinked band is observed, containing AHs (in green) of opposite orientations. The topology of the downstream TM and SH3 domains is indicated. Red triangles show the 3C cleavage sites. Scissors show the sites accessible to added protease. With the cysteine in the YG domain, three bands are observed, corresponding to the three topologies shown (see also ^14^).

Membranes were isolated from yeast cells and incubated first with the protease and then with an oxidant. When analyzed by reducing SDS-PAGE, two bands were seen (**Fig. 2a**), corresponding to the two orientations of the C terminus; the protease-resistant pool had the C terminus in the matrix, shielded by the membrane, as it could be cleaved only after membrane solubilization with detergent. Under oxidizing conditions, a single disulfide-linked band was observed after cleavage (**Fig. 2a**), which corresponds to an adduct of two PEX13 molecules with opposite orientations of their C termini. Analogous experiments with a cysteine introduced into the YG domain show three bands under oxidized conditions (**Fig. 2a**), as reported previously ^14^. These bands correspond to PEX13 dimers in which the two molecules have their C terminus either both in the matrix, on opposite sides of the membrane, or both in the cytosol (**Fig. 2b**). Thus, in contrast to crosslinked AHs, crosslinked YG domains consist of PEX13 molecules of both orientations. These experiments indicate that neighboring AHs have opposite membrane orientations and support the dual-membrane topology model in **Fig. 2b**.

The YG domain is required for proper channel formation. When the YG domain was deleted, PEX13 eluted in SEC as a broader peak at lower molecular weight than the wild-type protein (**Extended Data Fig. 5a**). A similar result was obtained when all tyrosines of the YG domain were mutated to serines (**Extended Data Fig. 5b**). Disulfide crosslinking with two cysteines introduced into this mutant demonstrated that the protein no longer forms a ladder and thus has a defect in oligomerization (**Extended Data Fig. 5c**). The channel is therefore stabilized by the YG phase.

### Cryo-EM structure of an inactive PEX13 channel

We noticed that the AH contains an unusual concentration of acidic amino acid residues (see sequence alignment in **Extended Data Fig. 6**). Two of these residues are conserved in most organisms, and additional ones in a subset of species. To test whether the acidic residues are required for peroxisomal protein import, we used an engineered pathway in *S. cerevisiae* cells that generates a pigment when import is impaired ^29^. Mutating only one of the three conserved acidic residues (E153, E167, E185) had no effect, but a double mutant (2E/A) showed reduced import activity and the triple mutant (3E/A) was essentially inactive (**Fig. 3a**). Mutating the equivalent three negatively charged residues in human PEX13 to alanines (E134A, D148A, D163A; 3ED/A mutant) did not affect the purification of the protein; like the wild-type protein, the mutant eluted in SEC as a sharp peak at high molecular weight (**Fig. 3b**). Together, these data suggest that the triple mutant forms an inactive channel.

**Fig. 3.**
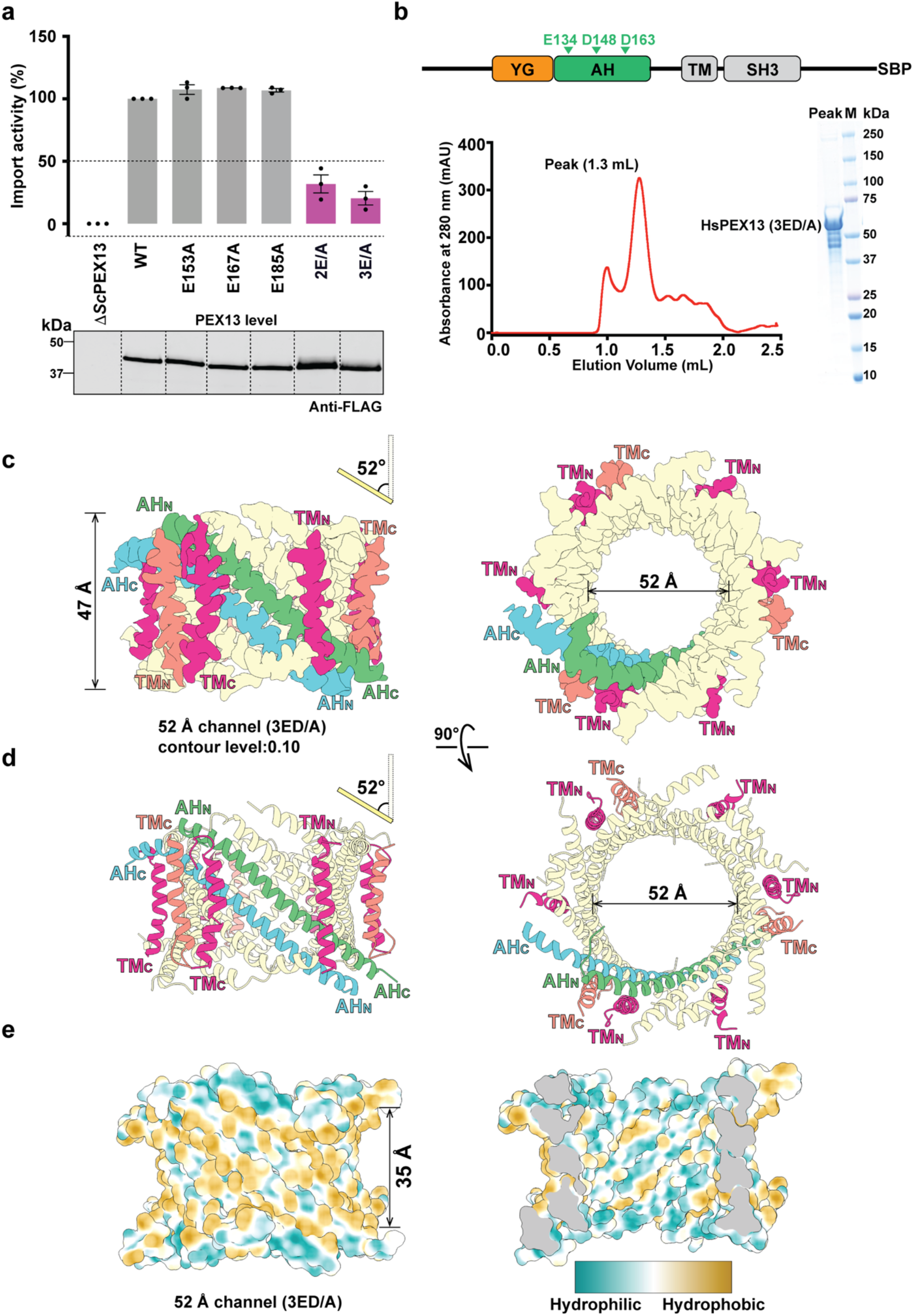
Cryo-EM structure of an inactive mutant channel. **a**, FLAG-tagged wild-type (WT) *S. cerevisiae* PEX13 or the indicated acidic residue mutants were expressed in *S. cerevisiae* cells lacking endogenous PEX13 (Δ*pex13*). Mutant 2E/A combines the E153A and E167A mutations, and 3E/A all three mutations. Peroxisomal protein import was measured with an engineered pathway that generates a pigment when import is impaired. Import activity is plotted relative to WT PEX13 (mean ± standard error of 3 experiments). The expression level of PEX13 is shown in the lower panel, analyzed by SDS-PAGE and immunoblotting with FLAG antibodies. **b**, Human PEX13 with three conserved acidic residues of the AH changed to alanines (3ED/A mutant) was expressed in *P. pastoris*. The protein was solubilized in digitonin, and purified with streptavidin beads followed by SEC (left panel). The peak fraction was analyzed by SDS-PAGE and Coomassie blue staining (right panel). **c**, Cryo-EM density map of the 3ED/A mutant channel in two views. Two AHs are highlighted in green and blue, with their N and C termini indicated. The helix tilt angle is shown on top of the left panel. Density for the TMs is colored salmon and red. **d**, Two views of the molecular model built into the map, shown as cartoons. **e**, Space-filling model showing the hydrophobic surface on the outside of the pore (left panel) and hydrophilic surface inside the pore (right panel). Hydrophobic and hydrophilic residues are colored with the scale on the right. The size of the hydrophobic belt is indicated.

The purified 3ED/A mutant was subjected to single-particle cryo-EM analysis. The side views in 2D averages showed that the heterogeneity of channel size was considerably reduced compared with the wild-type complex (**Extended Data Fig. 1d**). A map of 3.3 Å overall resolution could be obtained without imposing symmetry during refinement, but significant improvement was achieved after imposing 3-fold symmetry, resulting in a resolution of 2.8 Å (**Fig. 3c; Extended Data Table 1**). The final map shows a ring of 12 tilted AHs surrounding a central pore. The amino acid side-chain density was sufficient to model more than 60 residues of the AHs (**Fig. 3d; Extended Data Fig. 7a**). The tilt angle of the AHs (52°) and the pore diameter (∼52 Å) are similar to those of the small wild-type channel (53° and ∼52 Å, respectively). In contrast to the wild-type channels, which are formed from 6 copies of AH pairs, the mutant channel consists of 3 copies of four AH-containing units, a feature observed even without imposing symmetry during refinement. As before, the AHs alternate in their orientation around the ring, consistent with the dual topology in membranes and the placement of SH3 domains on both sides of the micelle (**Extended Data Fig. 1d**). The hydrophobic and hydrophilic faces of the AHs are again oriented outwards and inwards, respectively (**Fig. 3e**). The mutated negative charges all point into the pore (**Extended Data Fig. 8**). We speculate that the absence of charge repulsion in the 3ED/A pore disfavors channel widening.

Molecular models could also be built for 9 of the 12 TM segments (**Fig. 3c; d; Extended Data Fig. 7b**); the missing TM segments are likely flexible. The visible TM segments are positioned outside the pore and sit perpendicular to the membrane plane. They contact the outer, lipid-facing surface of the AH wall through hydrophobic residues. The interaction with a TM might be responsible for a kink of the AH near its N terminus. Some density seen in the unsharpened map inside the pore likely corresponds to the YG domains, arranged as sheets lining the hydrophilic faces of the AHs (**Extended Data Fig. 7c**).

### A large channel predicted by AlphaFold

The wild-type channel of ∼70 Å pore size is too small to allow the import of some cargo (see Discussion). However, AlphaFold2 predicts a large channel of ∼100 Å diameter with reasonable per-residue pLDDT scores and PAE plot (**Extended Data Fig. 4b**). The predicted channel contains highly tilted AHs (tilt angle ∼70°) (**Fig. 4a**). The C-terminal regions of the AHs (residues 177 to 199), which are invisible in the smaller pores, extend the helices to form the channel wall. The channel height is reduced to 29 Å (**Fig. 4a**). As with the smaller pores, the hydrophobic and hydrophilic sides of the AHs are oriented outward and inward, respectively, and the helix interfaces leave only small holes in the channel wall (**Fig. 4b**), giving further credence to the prediction.

**Fig. 4.**
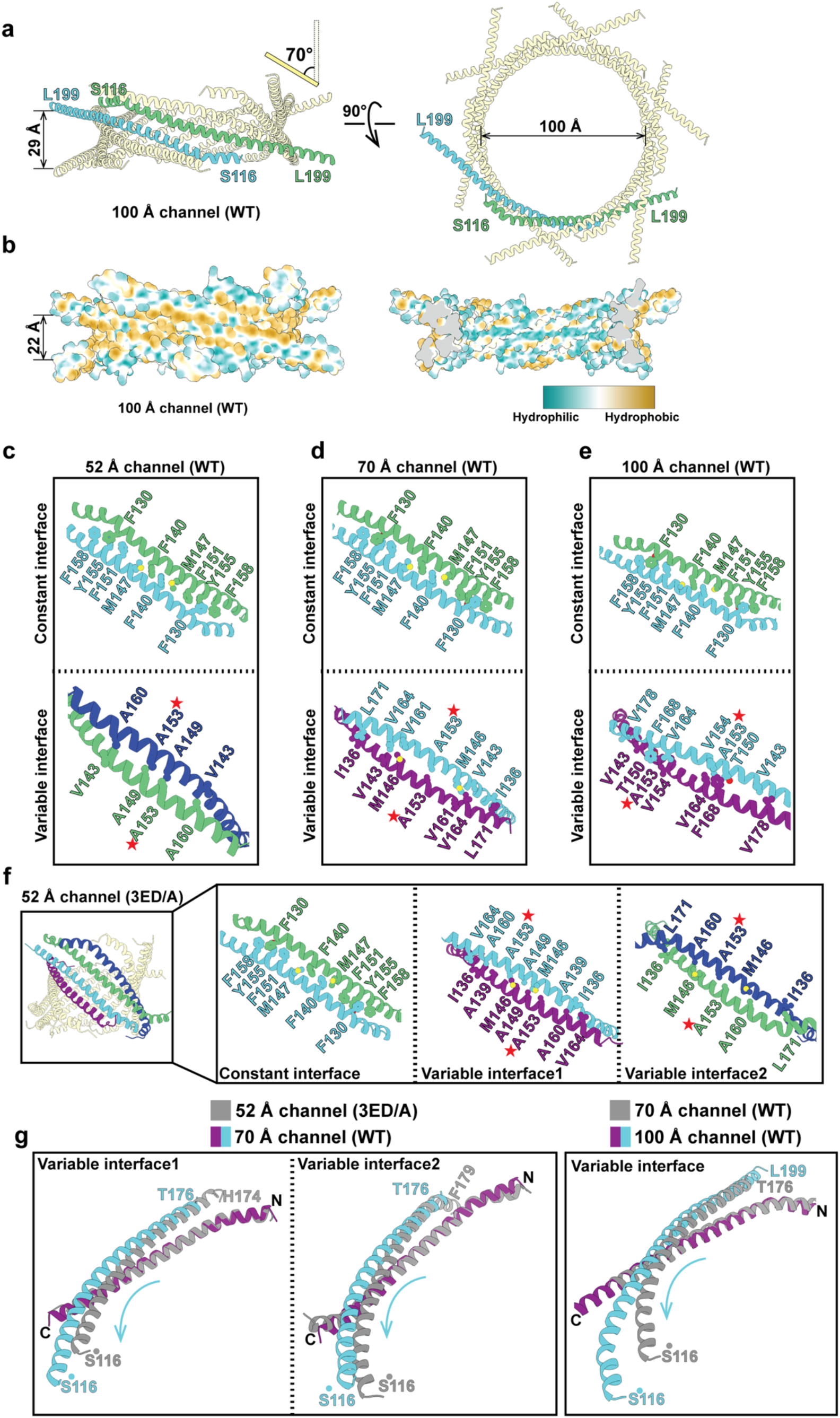
Helix sliding generates different pore sizes. **a**, Cartoon model of the AlphaFold2-predicted ∼100 Å channel in two views. Two AHs are highlighted. Channel height and helix tilt angle are indicated. **b**, Space-filling model of the channel, showing the hydrophobicity of the outside and inside of the pore (scale on the bottom). The size of the hydropobic belt is indicated. **c**, Shown are the constant and variable interfaces of neighboring AHs in the ∼52 Å wild-type (WT) channel. Interacting amino acids are labeled and shown with side chains. Hetero-atoms in side chains are shown in different colors. The position of the red star (residue 153) serves as a marker to show the relative movement of the helices. **d**, As in **c**, but for the ∼70Å channel. **e**, As in **c**, but for the ∼100 Å channel. **f**, As in **c**, but for the ∼52 Å 3ED/A mutant channel. This channel two distinct variable interfaces (1 and 2) on either side of the fixed AH dimer. Interface 2 is similar to the variable interface of the ∼52 Å wild-type channel. **f**, The left panels compare the variable interfaces of the ∼52 Å 3ED/A mutant channel with that of the ∼70 Å channel . One helix of the ∼52 Å channel (grey) was aligned with a helix of the ∼70 Å channel (purple). The movement of the other helix from the ∼52 Å channel (grey) to the ∼70 Å channel (blue) corresponds to an iris-like helix sliding. The right panel shows a comparison of the ∼70 Å and ∼100 Å channels.

Channels of different pore sizes are also predicted for *S. cerevisiae* PEX13. AlphaFold3 predicts channels of ∼52 Å and ∼70 Å, and AlphaFold2 a channel of ∼100 Å (**Extended Data Fig. 4c**). All predictions have low confidence scores, but the structures are similar to the corresponding predicted or observed channels of the human protein (**Extended Data Fig. 4a; b; Fig. 1e; f**). The predictions are further validated by the analysis of the interfaces of the AHs and crosslinking experiments (see below). Thus, peroxisomal import channels of different sizes seem to occur in evolutionarily distant organisms.

### An iris-like mechanism of pore size adjustment

The channel structures can be described as left-handed toroids. The left handedness is dictated by the chirality of the α-helices and the requirement that the hydrophobic sides face lipid. In all structures, each AH has two neighbors of opposite orientation. One of the two interfaces is constant and the other variable. The constant interface is the same in all channel structures of human PEX13 (**Fig. 4c-e**). It contains clusters of aromatic residues (Phe and Tyr) at the ends of both interacting helices (Phe130, Phe151, Tyr155, Phe158) and an aromatic residue at the center (Phe140). The aromatic residues lock the two helices together, so that they behave as a unit (fixed dimer). The amino acids in the constant interface are conserved between several different species (**Extended Data Fig. 6**).

The variable interface contains amino acids with small side chains, primarily Ala, Gly, Val, Thr and Ser (**Fig. 4c-e**). This interface allows neighboring helices to move relative to one another. Using residue A153 as reference (red star), the variable interface shifts by about one helix turn from the ∼52 Å to the ∼70 Å channel, and by additional ∼ 3 turns to the ∼100 Å channel (**Fig. 4c-e**). The 3ED/A mutant channel has two distinct variable interfaces on either side of the fixed dimer (interfaces 1 and 2), causing the observed 3-fold rotational symmetry (**Fig. 4f**). Interface 2 is almost identical to the single variable interface of the ∼52 Å wild-type channel (**Fig. 4c**). The comparison of these two channels shows that AH dimers can slide longitudinally along the helical axis with minimal changes of their tilt angle and pore diameter.

As in mammals, all three *S. cerevisiae* channels have the same constant interface with interacting aromatic residues and distinct variable interfaces containing small side-chain amino acids (**Extended Data Fig. 9a-c**). These similarities again support the validity of the AlphaFold predictions. As expected from the sequence features of the interfaces, mutation of three residues in the constant interface into alanines (Y170A, H174A, F177A, YHF/A mutant) , or conversion of one of the small residues (Ala172) in the variable interface to a bulky amino acid (Phe, A172F mutant) abolishes peroxisomal protein import in *S. cerevisiae* (**Extended Data Fig. 9d; e**).

The fixed dimers can undergo radial sliding along the variable interface to generate different tilt angles and corresponding pore sizes, as shown by superimposing these helices for the ∼52 Å and ∼70 Å channels, and for the ∼70 Å and ∼100 Å channels (**Fig. 4f**). As the pore widens from ∼52 Å to ∼100 Å, the rigid dimers slide past their neighbors around the ring, like the blades of a camera iris opening, moving farther from the pore’s center while tilting to lie flatter against the membrane (**Fig. 1e;f; Fig. 4a; Extended Data movie 1**). Whereas longitudinal sliding can occur with individual dimers, the radial change requires that all six dimers tilt in the same way. During sliding, the helices do not significantly rotate about their axis, thus maintaining the orientation of hydrophobic residues towards lipid. Channel widening is accompanied by a reduction of the membrane thickness surrounding the channel **(Extended Data movie 1**). The small and intermediate sized channels of ∼52 Å and ∼70 Å have hydrophobic belts of ∼32-35 Å thickness (**Fig. 1i; j**), which corresponds to the normal hydrophobic core of a phospholipid bilayer. The largest channel of ∼100 Å has a hydrophobic belt of ∼22 Å (**Fig. 4b**), which would cause distortion of the phospholipid bilayer. and may therefore have to be stabilized by receptor and cargo.

### Helix sliding confirmed by disulfide crosslinking

To verify that the sliding of the AHs generates different pores sizes, we performed disulfide crosslinking experiments with native *S. cerevisiae* membranes. FLAG-tagged PEX13 with single cysteines at different positions of the AH was expressed in yeast and isolated membranes were treated with an oxidant. The samples were analyzed by nonreducing or reducing SDS-PAGE (**Fig. 5a**). The observed crosslinks with a cysteine at position 160 can be explained with all three channel structures, as this residue is located at the constant interface and is in disulfide-bonding distance to the same residue in the neighboring helix (**Fig. 5b**). Crosslinks indicative of the variable interface in the ∼52 Å channel were observed with a cysteine at position 168 (**Fig. 5b**). Disulfide-bond formation with a cysteine at position 175 is consistent with the ∼70 Å and ∼100 Å channels, whereas crosslinks with position 180 can only be explained with the variable interface expected for the ∼100 Å channel (**Fig. 5b**); this residue would be too distant from the same residue in a neighboring helix in the other channel structures (>12 Å). Only little crosslinking was seen with several cysteine positions picked as controls (**Extended Data Fig. 10a**).

**Fig. 5.**
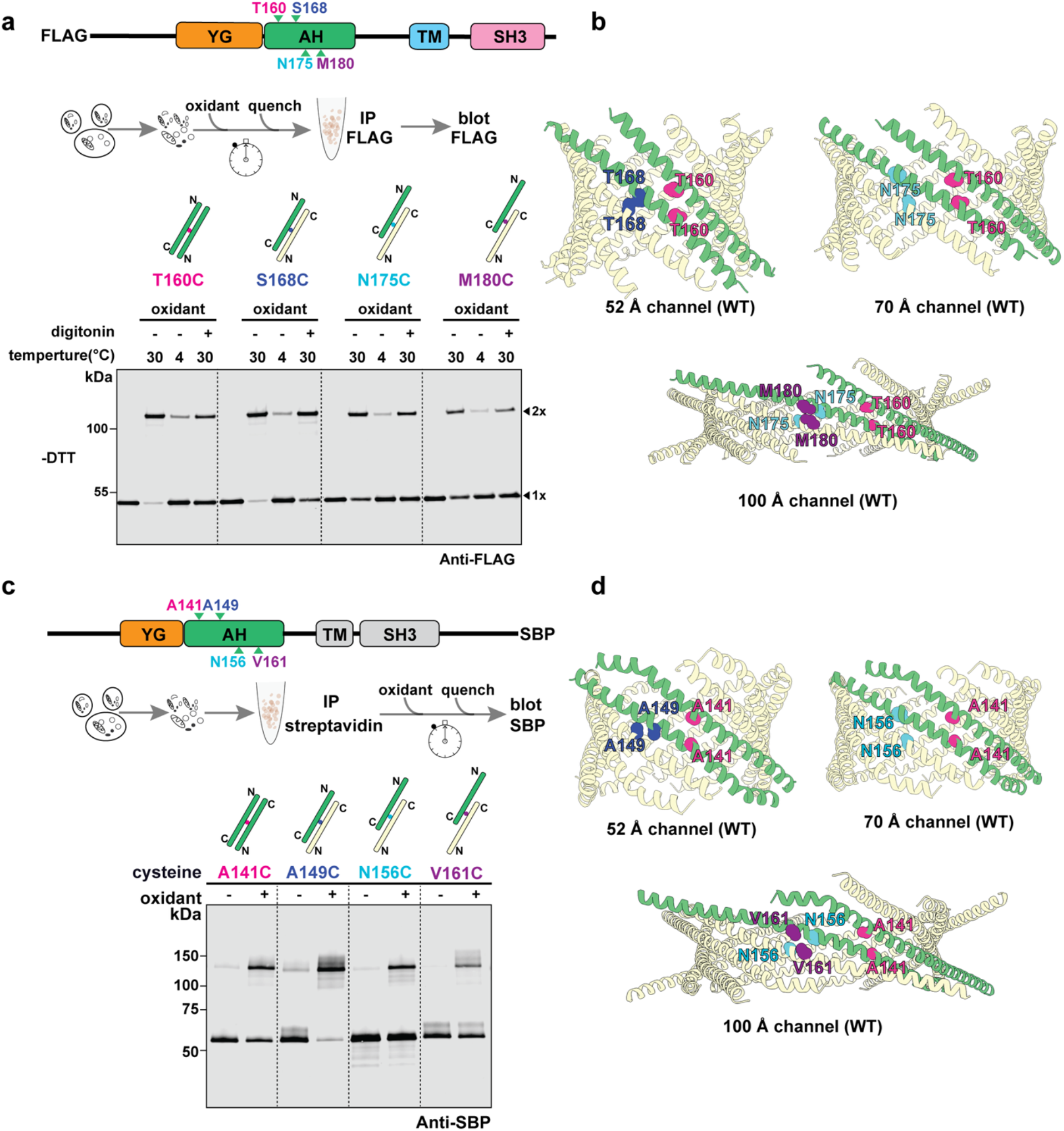
Different channel pores confirmed by disulfide crosslinking. **a**, FLAG-tagged PEX13 with cysteines at various positions of the AH was expressed in *S. cerevisiae*. Isolated membranes were treated with Cu-phenanthroline (oxidant) at 30°C or 4°C in the absence or presence of digitonin, as indicated. After quenching, the proteins were immunoprecipitated (IP) with FLAG antibodies and analyzed by nonreducing SDS-PAGE and immunoblotting with FLAG antibodies. Monomers (1x) and disulfide-linked dimers (2x) are indicated on the right. **b**, Cartoon models of the AlphaFold-predicted yeast channels. Two AHs connected by a constant interface are shown in green. Cysteines in neighboring AHs that explain disulfide-crosslinking in **a** are shown in the same color. Note that position 160 is in the constant interface; positions in the variable interface are diagnostic for the distinct channel states. **c**, SBP-tagged human PEX13 containing single cysteines at the indicated positions of the AH was expressed in *P. pastoris*. The proteins were solubilized in digitonin, purified with streptavidin beads, and treated with Cu-phenanthroline (oxidant) as indicated. The samples were analyzed by nonreducing SDS-PAGE and immunoblotting with SBP antibodies. **d**, As in **b**, but for the human channels.

**Fig. 6.**
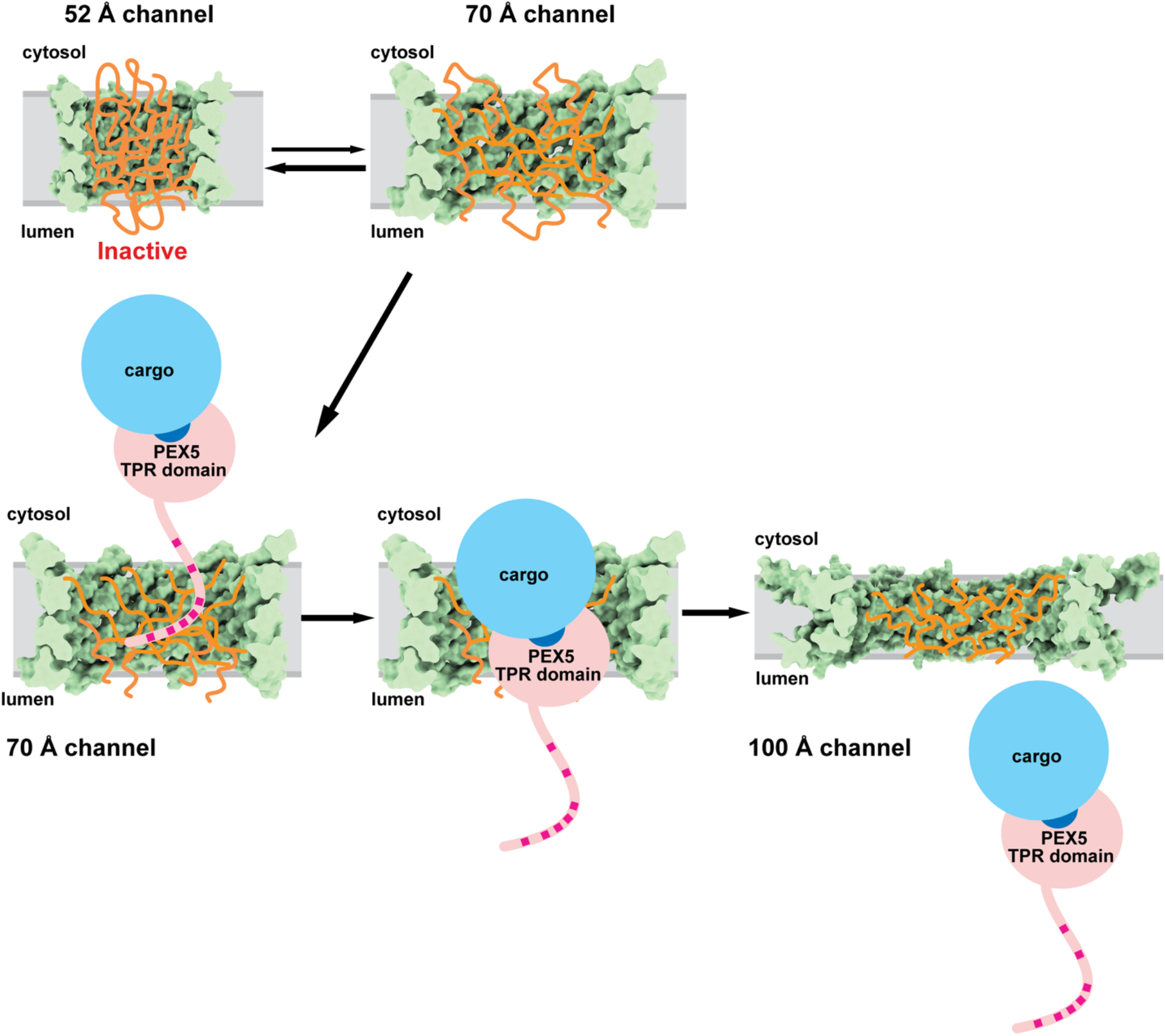
Protein import into peroxisomes reveals a novel translocation mechanism. Model of the cargo-size adaptable peroxisomal import channel. The different channels are shown as space-filling models in cut-away views. The YG meshwork is shown schematically in orange. The ∼52 Å pore corresponds to an inactive resting state that is in equilibrium with the ∼ 70 Å pore. The receptor PEX5 partitions into the YG phase through aromatic motifs in its unstructured N-terminus (red squares). Cargo containining a PTS1 sequence (dark blue) binds to the C-terminal TPR domain of PEX5. PTS2 cargo would additionally bind the adaptor PEX7 (not shown). The ∼70 Å pore can widen to ∼100 Å to allow peroxisomal import of large cargo.

Similar disulfide crosslinking experiments confirmed the different channels formed by purified human PEX13 (**Fig. 5c; d**). Position 141, located at the constant interface, is compatible with all three predicted channel structures. Crosslinks indicative of the variable interface in the ∼52 Å channel were observed with a cysteine at position 149 (**Fig. 5c; d**). Position 156 is consistent with the ∼70 Å and ∼100 Å channels; and position 161 is compatible only with the ∼100 Å channel. In both cases, a ladder was generated, although it was less pronounced with positions 141 and 161, suggesting that the ∼100Å channel is less abundant. (**Extended Data Fig. 10b**). Together, these data confirm the different channels generated by yeast and human PEX13.

## Discussion

Our structures show that the peroxisomal import channel consists of twelve PEX13 molecules. The walls consist of a ring of tilted AHs in alternating orientations, with hydrophobic faces on the outside, contacting the lipid, and hydrophilic faces inside, surrounding the YG phase. The channel can adopt different pore sizes by an iris-like movement of the AHs.

The pore of ∼52 Å is a resting state, into which the channel is locked when three conserved negative charges of the AH are mutated. This channel does not permit peroxisomal protein import in yeast, consistent with it being too narrow to allow passage of the C-terminal TPR domain of PEX5 (dimensions of ∼40 x 60 Å ^11^). Although peroxisomes are permeable to molecules smaller than ∼600 Da ^29,30^, a resting state with a dense YG meshwork inside the pore might be required to limit the exchange of metabolites between the cytosol and peroxisomes.

The ∼70 Å channel would be just large enough to accomodate the TPR domain ^31^ (**Extended Data Fig. 11a**). Monomeric matrix proteins bound to the TPR domain could also move through, as suggested by modeling the channel with a known structure of TPR-bound cargo in its interior ^31,32^ (**Extended Data Fig. 11b**). In fact, if the TPR domain and cargo passed the pore one after the other, the ∼70 Å pore could transport even the largest known monomeric matrix protein (∼79 kDa ^33^, corresponding to a spherical diameter of 57 Å), as well as oligomeric proteins known to be imported as monomers (72-75 kDa) ^34–36^. Monomeric PTS2 cargoes bound to the adaptor PEX7 (32 x 45 Å) and a short receptor segment could also move through the ∼70 Å channel. The YG domains would likely have to be displaced, but a similar situation is seen in the nuclear import of the HIV-1 capsid, which interacts with the FG repeats, but has a diameter that is almost identical to the width of the nuclear pore ^37,38^.

The ∼52 Å channel is likely the most stable state but may be in equilibrium with the ∼70 Å channel, as both structures co-exist in the cryo-EM dataset. The spontaneous switching between the two states may be facilitated by both channels sitting in an undistorted lipid bilayer. Escape from the ∼52 Å channel state requires the conserved acidic residues in the AH; they all point into the pore and are most tightly packed in the ∼52 Å channel (**Extended Data Fig. 8**), possibly generating electrostatic repulsion that could facilitate channel widening.

Although the ∼70 Å channel might be sufficient for some cargo, channel widening is certainly required for others. The predicted ∼100 Å channel could accommodate the largest oligomeric proteins, such as the tetramer of isocitrate lyase (∼256 kDa) ^39^. It would also be needed for the observed import of 9 nm gold particles coated with PTS1 peptides ^40^. Although an experimental structure of the ∼100 Å channel is currently unavailable, the AlphaFold-prediction is supported by (1) the hydrophobic and hydrophilic surfaces of the AHs pointing outwards and inwards, respectively, (2) the channel walls having only small holes; (3) the AHs sharing the constant interface with the experimentally observed channels, and (4) the variable interface being independently confirmed by disulfide crosslinking with intact membranes and purified protein. The ∼100 Å channel is probably energetically unfavorable, as it would sit in a thinned and distorted lipid bilayer and has weaker AH interactions than the smaller channels. Although challenging, it will be important to test experimentally whether channel widening is induced when PEX5 and large cargo move through.

Although we have not directly visualized the transitions between the different channel states, they likely occur by a simple iris-like motion of the AHs. Two neighboring AHs of opposite orientation form a fixed dimer that can slide radially relative to the neighboring dimer, thus generating different helix tilt angles and pore sizes (**Fig. 4f; Extended Data movie 1**). Switching from one pore size to the other requires that all variable helix interfaces change. A fixed dimer can also slide longitudinally relative to its neighbor. This movement explains the two distinct variable interfaces in the mutant channel and results in only minimal changes of the pore size. Longitudinal sliding might explain the low resolution of the wild-type protein structures.

The partitioning of the receptor/cargo complex into the YG phase could shift the equilibrium from the resting state (∼52 Å pore) to the active pores, possibly by expansion of the volume of the YG phase, which in turn could push the walls outwards. Oligomeric cargoes can have more than one bound receptor molecule, so their simultaneous partitioning into the YG phase could cause the conversion into the ∼100 Å channel. Whether the channel adopts only discrete pore sizes (∼52 Å, ∼70 Å, ∼100 Å) is unclear. Intermediates are conceivable, as the small side-chain amino acids make the variable interface promiscuous, probably allowing additional arrangements of the fixed dimers. Specifically, the change of the pore size from ∼70 Å to ∼100 Å is substantial, suggesting that intermediates may be required for this transition.

The iris-like changes of the channel pore are based on the unusual dual membrane topology of PEX13. This feature creates the constant and variable interfaces that enable helix sliding, and allows the YG phase to be suspended inside the membrane and be insulated from the surrounding lipid bilayer. It also places the SH3 domains of PEX13 alternately on either side of the membrane. SH3 domains facing the cytosol could recruit PEX5 to the channel by binding to the receptor’s aromatic motifs ^41,42^. In yeast, SH3 domains facing the matrix could recruit PEX14 ^43^ but in mammals, PEX14 recruitment is mediated by different interactions ^41^.

Our results support a new mechanism of protein translocation, distinct from previously described mechanisms that use either an aqueous channel or a locally thinned membrane to lower the energy barrier for moving a polypeptide across the membrane. An aqueous channel is formed by the prototypical Sec61 complex in the endoplasmic reticulum (ER) membrane and the analogous SecY complex in the plasma membrane of prokaryotes (for review, see ^44^). Aqueous channels are also used for translocation across the outer membranes of mitochondria and chloroplasts ^45,46^. In these cases, only unfolded polypeptides can move through the narrow channels. The locally thinned membrane mechanism is exemplified by bacterial YidC, which serves as an insertase of membrane proteins ^47,48^, and the Hrd1 complex ^49^, which is responsible for moving misfolded ER proteins into the cytosol for proteasomal degradation (ERAD). This mechanism is also used in other translocation systems, including that in the inner mitochondrial membrane (for review, see ^50^). By contrast, our results suggest that the peroxisomal import channel has a unique cargo-adaptable pore that could allow folded and even oligomeric cargo to pass through ^51^. The lining of the channel walls with amphipathic helices is also peculiar, found to our knowledge only in α-helical pore-forming toxins, whose pores are much smaller pores ^52,53^.

The peroxisomal import channel resembles the nuclear pore. Both are formed from oligomeric assemblies of components with rotational symmetry, twelve in the case of PEX13 and multiples of eight in the case of nucleoporins. Like the peroxisomal channel, the nuclear pore contains a selective phase ^16^ and can probably adjust to cargo size ^37,38,54^. However, an important distinction is that the peroxisomal YG phase is positioned within the membrane, whereas the FG phase in nuclear pores is located in the aqueous space that connects cytosol and nucleoplasm. Nevertheless, the similarities between peroxisomal import and nuclear transport suggest common solutions to the challenge of transporting folded proteins between compartments.

## Acknowledgments

We thank Jeremi Kuklewicz for insightful discussions and other members of the lab for helpful comments. We thank Vlad Denic, Jeremi Kuklewicz, David Pellman, and Mike Skowyra for critical reading of the manuscript. We thank Chad Hicks for help with preliminary cryo-EM data collection at the Boston University Medical Campus facility, which is supported by NIH grant S10-OD032253. We also thank Shixin Yang and Zhening Zhang for cryo-EM data collection at the HHMI Janelia Research Campus Cryo-EM Facility. This article is subject to HHMI’s Open Access to Publications policy. HHMI lab heads have previously granted a nonexclusive CC BY 4.0 license to the public and a sublicensable license to HHMI in their research articles. Pursuant to those licenses, the author-accepted manuscript of this article can be made freely available under a CC BY 4.0 license immediately upon publication. This manuscript is the result of funding by the National Institutes of Health (NIH). It is subject to the NIH Public Access Policy. Through acceptance of this federal funding, NIH has been given a right to make this manuscript publicly available in PubMed Central upon the Official Date of Publication.

## Funding

National Institutes of Health grant R01GMO52586 (TAR)

Howard Hughes Medical Institute (HHMI) (TAR)

## Author contributions

T.A.R., D.W.H., and Y.G. conceptualized the study. D.W.H. designed the strategies for protein expression, purification, mutagenesis, and structure determination. D.W.H. and Y.G. performed the crosslinking experiments. T.A.R. supervised the project and wrote the original draft.

## Data and materials availability

All data are available in the main text or the supplementary materials. Unique reagents generated by this study are available from the lead contact, Tom A. Rapoport with a completed Materials Transfer Agreement. Cryo-EM maps and atomic coordinates generated in this work are available from the Electron Microscopy Data Bank (EMDB; EMD-78756, HsPEX13 52Å import channel (WT); EMD-78762, HsPEX13 70Å import channel (excluding 177-186); EMD-78763, HSPEX13 70Å import channel (including 177-186)). The accession codes for the HsPEX13 52Å import channel (3ED/A) are EMD-78822 in EMDB and PDB:38HQ in the Protein Data Bank.

## Supplementary Materials

### Materials and Methods

All reagents were obtained from Millipore-Sigma unless specified otherwise. Buffers were prepared in ultrapure water. RT denotes room temperature.

#### Plasmids for expression of human PEX13 (HsPEX13) in *Pichia pastoris*

For overexpression of *HsPEX13* in *P. pastoris*, the DNA sequence encoding *HsPEX13* (UniProt accession no. Q92968) was synthesized and codon-optimized by Integrated DNA Technologies. *HsPEX13* carried at the N terminus three FLAG epitopes followed by a TEV protease cleavage site. The C terminus was fused to a streptavidin-binding protein (SBP) tag preceded by an HRV 3C protease cleavage site. All segments were assembled into the integrating vector pPICZα-LINK using Gibson assembly. For expression of the *HsPEX13*(3ED/A) mutant in *P. pastoris*, residues E134, D148, and D163 were mutated to alanines. For expression of the *HsPEX13* ΔYG (1–110) mutant, residues 1–110 of the YG domain were deleted. For disulfide crosslinking experiments, the residues indicated in the text were changed to cysteines. In these mutants, the endogenous cysteine residue at position 220 was mutated to serine (C220S). All deletions and single-residue mutants were generated by PCR or site-specific mutagenesis. For the HsPEX13 SG mutant, the tyrosine residues in the YG domain of *HsPEX13* (Y75, Y82, Y85, Y90, Y93, Y96, Y98, Y100, and Y105) were replaced with serines, cysteines were introduced at positions 79 and 101, and the endogenous cysteine at position 220 was changed to serine by *de novo* gene synthesis (Integrated DNA Technologies). All constructs were verified by whole-plasmid DNA sequencing.

#### Expression of HsPEX13 in *Pichia pastoris* and protein purification

*P. pastoris* wild-type strain SMD1168 was transformed by electroporation. Transformed yeast cells were grown on Zeocin-containing YPDS plates (YPD medium supplemented with sorbitol) at 30°C for 3–4 days. A single colony was picked to inoculate a starter culture, which was incubated overnight at 30°C. A large-scale culture was then inoculated by diluting the starter culture 1:80 into buffered minimal glycerol (BMG) medium. The culture was incubated at 30°C for approximately 24 h, after which protein expression was induced by transferring the cells into an equal volume of buffered minimal methanol (BMM) medium. After an additional 18–20 h of incubation at 28 °C, the cells were harvested by centrifugation at 4,500 × *g* for 15 min at 4°C. The cell pellets were flash-frozen and stored at −80°C until further use.

For purification of wild-type *HsPEX13, the 3ED/A mutant, the* ΔYG mutant, and the SG mutant, ∼10-15 g of cell pellets were resuspended in 80 ml of buffer A (400 mM NaCl and 25 mM HEPES, pH 7.5) supplemented with 1 mM phenylmethanesulfonyl fluoride (PMSF). Cells were lysed using a BioSpec BeadBeater for 50 min with 20 s on/60 s off cycles in a water–ice bath. The homogenate was centrifuged at 4,000 × *g* for 20 min to remove cell debris. The supernatant was then subjected to ultracentrifugation in a Ti45 rotor (Beckman Coulter) at 42,000 r.p.m. for 1 h at 4 °C. The membrane pellet was resuspended using a Dounce homogenizer in 20 ml of buffer A supplemented with 1% digitonin (EMD Millipore) and a protease inhibitor cocktail (Bimake, no. B14001), followed by incubation for 60 min at 4°C. Insoluble material was removed by centrifugation in a Ti70 rotor (Beckman Coulter) at 42,000 r.p.m. for 30 min. The supernatant was transferred to a new tube and incubated with 1 ml of high-capacity streptavidin resin (Thermo Scientific) for 1.5 h. The resin was washed with 20 ml of buffer A containing 0.1% digitonin, and the bound protein was eluted with 5 ml of buffer B (150 mM NaCl, 25 mM HEPES, pH 7.5, 10% glycerol and 0.1% digitonin) supplemented with 2 mM biotin (Sigma).The eluted protein was concentrated using a 100-kDa molecular-weight-cutoff filter (Millipore) and further purified by size-exclusion chromatography on a Superose 6 Increase 3.2/300 column (Cytiva Life Sciences), equilibrated with buffer C (25 mM HEPES, pH 7.5, 150 mM NaCl and 0.06% digitonin).

#### Disulfide-crosslinking of HsPEX13 variants

Single-cysteine mutant proteins of HsPEX13 were expressed in *P. pastoris* as above. The cells (0.1-0.5 g) were collected by centrifugation, washed once with water, and resuspended in ice-cold lysis buffer containing 300 mM NaCl and 25 mM HEPES (pH 7.5), supplemented with 1 mM PMSF and a protease inhibitor cocktail (Bimake, no. B14001). Each cell suspension was transferred to 2-ml screw-cap tubes (Sarstedt, no. 726994005) and kept on ice. Pre-chilled 0.5-mm glass beads were added, and the cells were lysed using a Mini-BeadBeater-16 (BioSpec Products, no. 607) in four 30-s cycles, with 2-min cooling intervals on ice in between. The lysate was clarified by centrifugation at 2,000 × g for 5 min at 4°C to remove intact cells and cell debris. The resulting supernatant was further centrifuged at 20,000 × g for 10 min at 4°C to pellet heavy membranes, including peroxisomes. The pellet was gently resuspended in lysis buffer supplemented with 1% digitonin by pipetting up and down, and then incubated on a rotator for 1 h at 4°C. Insoluble material was removed by centrifugation at 20,000 × g for 10 min at 4°C. The resulting supernatant was incubated with high-capacity streptavidin resin (Thermo Scientific) for 1 h at 4°C. The resin was washed with lysis buffer containing 0.1% digitonin, and the bound proteins were eluted with buffer B containing 150 mM NaCl, 25 mM HEPES (pH 7.5), 10% glycerol, and 0.1% digitonin, supplemented with 2 mM biotin. The eluted proteins were immediately snap-frozen and stored at −80°C until further use.

The samples were thawed and added to assay buffer containing 0.1% digitonin, 20 mM HEPES–NaOH (pH 7.5), 50 mM NaCl, 250 mM sucrose, and 1 mM EDTA, and then treated with the oxidizing agent copper(II)-phenanthroline at 0, 0.3 mM, or 0.5 mM for cysteines in the AH, and at 0, 0.05 mM, or 0.2 mM for cysteines in the YG domain. The samples were incubated for 30 min at 30°C with gentle agitation, quenched with 10 mM N-ethylmaleimide (NEM). One half of the sample received 100 mM dithiothreitol (DTT) and both were heated at 95°C in Laemmli buffer for SDS-PAGE and immunoblotting.

For disulfide crosslinking with two cysteines incorporated into HsPEX13, the following mutants were used: S79C, N101C, C220S; A141C, A149C, C220S; A141C, N156C,C220S; A141C, V161C,C220S; and A141C, A149C, C220S, E134A, D148A, D163A. These proteins were purified as above, except that they were further purified by SEC on a Superose 6 Increase column. The peak fractions were used for disulfide crosslinking as described before.

#### Peroxisomal import assay in *S. cerevisiae* cells

A modified violacein biosynthetic pathway ^27^ was used to quantitatively measure peroxisomal matrix protein import in *S. cerevisiae* cells, as described ^11^. In this system, the first two enzymes of the pathway (VioA and VioB) reside in the cytosol, whereas the third enzyme (VioE) contains a peroxisome targeting signal and is sequestered inside peroxisomes when import is functional. Only when import is compromised does VioE accumulate in the cytosol and the green pigment is made. Pigment production is inversely proportional to import efficiency ^27^.

Import was measured in a strain (UTL7A (*MATa, ura3-52, trp1, leu2-3/112*)) that contains an expression cassette encoding the three enzymes (VioA, VioB, and VioE) integrated into the genome along with a leucine auxotrophic marker. The *pex13* gene was deleted and replaced with the *natMX* marker (nourseothricin resistance), as described ^11^. This strain was transformed with PCR-generated DNA fragments coding either for wild-type PEX13 or for PEX13 mutants carrying changes of the negatively charged residues (E153, E167, E185) to alainines, all with 60-bp homology overhangs. The PCR products were introduced into yeast cells by lithium acetate–based transformation ^48^. Transformants were selected on YPD medium containing hygromycin (250 μg/ml; Fisher scientific). Correct integration of the mutant or epitope-tagged coding sequence was validated by PCR and DNA sequencing.

For pigment quantification, yeast strains were cultured overnight in YPD medium at 30°C with shaking. The next morning, cultures were diluted 150-fold into 3 ml of freshly prepared synthetic defined (SD) medium lacking leucine and incubated at 30°C with shaking for an additional 50 h. Cells were collected by centrifugation and resuspended in 300 µl of glacial acetic acid (Fisher Scientific). The cell suspensions were transferred to 1.5-ml microcentrifuge tubes, incubated at 95°C for 10 min, mixed by inversion, and incubated for an additional 10 min at 95°C. Cell debris were removed by centrifugation at 13,000 × *g* for 5 min at room temperature. Subsequently, 200 µl of each supernatant was transferred to a 96-well, round-bottom, black-walled plate (Corning, no. 3792). Fluorescence was measured using a BioTek Synergy Neo2 microplate reader with excitation and emission wavelengths of 535 ± 10 nm and 585 ± 10 nm, respectively. Relative peroxisomal matrix protein import activity was calculated by normalizing the fluorescence signal of each strain to the average fluorescence of wild-type cells (set to 100%) and *pex13Δ* cells (set to 0%), using linear interpolation.

To assess the expression levels of the FLAG-tagged PEX13 constructs, membranes from the relevant strains were solubilized in 1.0% (w/v) dodecylmaltoside (DDM), and subjected to immunoprecipitation using anti-FLAG M2 beads before analysis by SDS-PAGE.

To evaluate the functional significance of the constant and variable AH interfaces, Y170, H174, and F177 at the constant interface were mutated to alanines, and A172 at the variable interface was mutated to phenylalanine. All constructs encoding the PEX13 variants were transformed into yeast cells, and peroxisomal import activity was measured as described before. To assess protein expression levels, all PEX13 constructs carried a FLAG tag. Expression levels were determined by SDS-PAGE followed by immunoblotting with FLAG antibodies.

#### Disulfide-crosslinking of *S. cerevisiae* PEX13

The Δ*pex13* strain described above was transformed with PCR products coding for FLAG-tagged PEX13 variants containing the indicated cysteine substitutions to integrate the genes into the genome. The cells were cultured overnight in YPD medium at 30°C, collected by centrifugation, washed once with water, and resuspended in ice-cold lysis buffer containing 20 mM HEPES–KOH (pH 6.8 at room temperature), 150 mM potassium acetate, 5 mM magnesium acetate, 250 mM sorbitol, and 1 mM EDTA, supplemented with 1 mM PMSF and a protease inhibitor cocktail (Bimake, no. B14001). The cell suspension was transferred to 2-ml screw-cap tubes (Sarstedt, no. 726994005) and kept on ice. Prechilled 0.5-mm glass beads were added, and cells were lysed using a Mini-BeadBeater-16 (BioSpec Products) for four 30-s cycles, with 2-min cooling intervals on ice between cycles. The lysate was clarified by centrifugation at 2,000 × g for 5 min at 4°C to remove intact cells and cell debris. The resulting supernatant was centrifuged at 20,000 × g for 10 min at 4°C to pellet peroxisomes. The final pellet was gently resuspended in assay buffer containing 20 mM HEPES–NaOH (pH 7.2), 50 mM NaCl, 250 mM sucrose, and 1 mM EDTA, followed by incubation on a rotator at 4°C for 15 min.

The homogeneous membrane suspension was treated with or without the oxidizing agent copper (II)-phenanthroline (Cu-phenanthroline) at 0.1 mM for 30 min at 30°C or 4°C with gentle agitation. Where indicated, 1.0 % digitonin was added before the oxidant to solubilize the membranes. Unreacted cysteine residues were quenched with 10 mM NEM at 30°C for 5 min. PEX13 was immunoprecipitated from the resulting supernatant using anti-FLAG M2 agarose beads (Millipore, no. A2220) and analyzed by SDS-PAGE.

#### Determination of the membrane topology of the AH in *S. cerevisiae*

A single cysteine was introduced at position 168 (AH) into PEX13 containing a FLAG tag at its N-terminus and a 3C protease-cleavage site and SBP tag at the C terminus. The construct was integrated into the chromosome. An analogous strain with a cysteine at position 104 (YG domain) has been described before ^11^. Membranes from these strains were prepared as described above, mixed with 2 μM homemade 3C protease, and incubated for 16 h at 4°C. The samples were then treated with copper(II)-phenanthroline (0.1 mM) for 30 min at 30°C to promote disulfide bond formation. The reactions were subsequently quenched with 10 mM NEM and the membranes were solubilized with 0.5% dodecylmaltoside (DDM) for 1 h at 4°C with shaking. PEX13 was then immunoprecipitated from the resulting supernatant using anti-FLAG M2 agarose. The precipitated material was eluted with 0.4 mg/mL 3×FLAG peptide (Bimake, no. B23112) in assay buffer containing 0.05% (w/v) DDM and prepared for reducing or non-reducing SDS-PAGE.

#### Electrophoresis and immunoblotting

Samples were analyzed by SDS-PAGE on 4–20% TGX precast polyacrylamide gels (Bio-Rad). For immunoblotting, proteins were transferred onto nitrocellulose membranes (Bio-Rad, no. 1620112) using Trans-Blot turbo Transfer System (BioRad). Membranes were blocked in TBST buffer (20 mM Tris–NaOH, pH 7.5, 150 mM NaCl, and 0.1% (v/v) Tween 20) containing 3% (w/v) non-fat milk solids (Apex, no. 20241) and then incubated with antibodies against the FLAG epitope (Millipore Sigma, no. F7425) or SBP tag (Millipore Sigma, no. MAB10764), followed by fluorescently labeled secondary antibodies IRDye 800CW (LI-COR Biosciences), as appropriate. Blots were imaged using a LI-COR Odyssey M imaging system.

#### Cryo-EM sample preparation and data acquisition

For cryo-EM sample preparation of wild-type *HsPEX13,* the peak fractions obtained from SEC were concentrated to ∼2.6 mg/ml. Three μl of the sample were applied to Quantifoil Cu 1.2/1.3 400 mesh grids that were glow-discharged at 15 mA for 30 s using a PELCO glow-discharge system (Ted Pella). The grids were blotted for 7.0 s at a blot force of 20 and ∼100% humidity and then plunge-frozen in liquid ethane using a Vitrobot Mark IV instrument (Thermo Fisher Scientific). For cryo-EM sample preparation of *HsPEX13* (3ED/A), the SEC peak fractions were concentrated to ∼5.0 mg/ml. Three μl of the sample were applied to a glow-discharged Quantifoil grid (1.2/1.3, Au, 300 mesh). The grids were blotted for 6.0 s at a blot force of 6 and ∼100% humidity and then plunge-frozen in liquid ethane using a Vitrobot Mark IV instrument. All grids were stored in liquid N_2._

Cryo-EM data were collected on a Titan Krios electron microscope (FEI) operated at 300 kV and equipped with a post-GIF Gatan Summit direct electron detector K3 camera (Gatan) at the HHMI Janelia Research Campus Cryo-EM Facility. For wild-type *HsPEX13,* images were collected on Krios2. An energy-filter slit width of 20 eV was used to remove inelastically scattered electrons. All cryo-EM movies were recorded in super-resolution counting mode using SerialEM. A nominal magnification of ×81,000 corresponded to a calibrated physical pixel size of 1.06 Å per pixel and a super-resolution pixel size of 0.53 Å per pixel. The total electron dose was 50 electrons Å⁻², fractionated over 50 frames. The defocus range was set from −1.0 to −2.5 μm. A total of 9,697 movies were recorded over 3 three days.

For the *HsPEX13* (3ED/A), images were collected on Krios1. All cryo-EM movies were recorded in super-resolution counting mode using SerialEM. A nominal magnification of ×81,000 corresponded to a calibrated physical pixel size of 0.857 Å per pixel and a super-resolution pixel size of 0.4285 Å per pixel. The defocus range was set from −1.0 to −1.8 μm. A total of 17,301 movies were recorded over 3 days. All other data-collection parameters were the same as those used for the wild-type dataset.

#### Data processing

For wild-type (WT) HsPEX13, cryo-EM data were processed as summarized in **fig. S2**. Data processing was performed using cryoSPARC v4.7.1 ^49^. Following patch motion correction, patch-based contrast transfer function (CTF) estimation, and manual inspection to remove micrographs with ice contamination or poor image quality, particles were picked from a random subset of 500 micrographs using the cryoSPARC Blob Picker. Two-dimensional (2D) classification was then performed to generate 2D templates, and particles were autopicked from 8,712 micrographs using these templates with the cryoSPARC Template Picker and Inspect Particles, yielding a total of 1,561,361 particles. Particles were initially extracted into 180 × 180-pixel boxes with 2× downscaling and subjected to two rounds of reference-free 2D classification. Poorly populated 2D classes were removed, yielding a selected set of 650,685 particles. These particles were then re-extracted into 360 × 360-pixel boxes and subjected to *ab initio* reconstruction (N=6) to generate initial 3D models. *Ab initio* reconstruction was performed with a maximum resolution of 6 Å, an initial resolution of 9 Å, an initial minibatch size of 300, a final minibatch size of 1,000, and C6 symmetry. Two distinct reconstruction classes were obtained, corresponding to large (∼70 Å channel)- and small (∼52 Å channel)-size particles, containing 172,462 and 135,666 particles, respectively.

For the ∼70 Å channel reconstruction, two distinct classes were obtained after two rounds of ab-initio reconstruction (N=3). One reconstruction lacked density corresponding to the tail region (residues K177–L186) and contained 50,745 particles, whereas the other contained 49,051 particles. Each particle subset was subjected to non-uniform refinement ^50^ with C6 symmetry, yielding global resolutions of 6.18 Å and 6.23 Å, respectively.

For the small-size reconstruction, 36,627 particles from the cleanest 3D class were selected after one round of ab-initio reconstruction (N=5) and subjected to non-uniform refinement with C6 symmetry, yielding a global resolution of 6.2 Å. Otherwise, the parameters used for ab initio reconstruction were the same as above.

The data processing for the HsPEX13 (3ED/A) dataset is summarized in **Extended Data Fig. 2**. Data processing was performed using cryoSPARC v4.7.0 ^49^, as described above, with particles picked from a random subset of 3,517 micrographs by cryoSPARC Blob Picker. Two-dimensional (2D) classification was then performed to generate 2D templates, and particles were autopicked from 15,130 micrographs using these templates with the cryoSPARC Template Picker and Inspect Particles, yielding a total of 3,213,428 particles. Particles were initially extracted into 180 × 180-pixel boxes and subjected to two rounds of reference-free 2D classification. Poorly populated 2D classes were removed, yielding a selected set of 704,254 particles. These particles were then re-extracted into 360 × 360-pixel boxes and subjected to an additional round of 2D classification to yield 586,960 particles. The resulting particle set was subjected to two rounds of ab-initio reconstruction (N = 6 in the first round and N = 3 in the second round) to generate initial 3D models. Ab-initio reconstruction was performed with a maximum resolution of 6 Å, an initial resolution of 9 Å, an initial minibatch size of 300, a final minibatch size of 1,000, and C3 symmetry. One reconstruction class was selected, containing 102,845 particles. This particle set was subsequently subjected to one round of heterogeneous refinement (N=3) using a 360-pixel refinement box and C3 symmetry, yielding a final particle set of 75,550 particles. The resulting particle set was subjected to non-uniform refinement with C1 or C3 symmetry, resulting in global resolutions of 3.3 Å and 2.8 Å repectively. Histograms of directional Fourier shell correlation curves and sphericity values were calculated with the 3DFSC server ^51^. All software used in this study was supported by SBGrid ^52^.

#### Model building

Wild-type HsPEX13 models were derived from AlphaFold-predicted models. For the 52 Å model, the amphipathic helix (S116–L171) was fitted into the small-size density map using ChimeraX 1.8 ^53^. TM helices and portions of the AH that extended beyond the density were removed in Coot ^54^.

For the ∼70 Å structure without the tail region, the AH (S116–T176) was fitted into the density map, and portions of the model that extended beyond the density were trimmed in Coot. For the 70 Å model containing the tail region (K177–R186), the AlphaFold model was first fitted into the density map, followed by manual adjustment of residues K177–R186 to better fit the corresponding density. Additional regions of the AlphaFold model that were not supported by the density were removed.

For the 52 Å channel of *HsPEX13* (3ED/A), the small-size AlphaFold model was docked into the density map and subsequently transferred to Coot for manual model building. The model was refined through iterative rounds of automated refinement in Phenix ^55^. Local resolution was estimated using cryoSPARC ^49^. Structural visualization and analysis were performed using ChimeraX 1.8 ^53^.

To model PEX5 inside the ∼70 Å channel, the TPR domain of HsPEX5 (PDB: 9GAG) was manually docked into the channel. Modeling of receptor–cargo complexes inside the channel was done with ScPEX5 TPR domain associated with Eci1 (PDB: 9FH0) and with HsPEX5 TPR domain associated with mSCP2 (PDB: 2C0L). Docking was performed with ChimeraX 1.8.

#### AlphaFold predictions of PEX13 structures

The ∼52 Å, ∼80 Å and ∼70 Å channel models of *HsPEX13* and *Sc*PEX13 were predicted using AlphaFold3 through the AlphaFold Server (https://alphafoldserver.com/) The ∼100 Å channel models of *HsPEX13* and *Sc*PEX13 were predicted using AlphaFold2 ColabFold through the COSMIC2 server (https://www.cosmic2.org/).

In initial predictions, we used 12 copies of the AH of HsPEX13, spanning residues S116–G200, and identified channel states with approximately 52 Å ,70 Å, and 80 Å in diameter. However, the prediction confidence scores were low. We found significant improvement of the predictions when *12* copies of AHs, comprising residues S116–G200, were supplemented with 12 copies of the TM, comprising residues A227–T256. In this case, only channels of ∼52Å, and ∼70Å diameter were predicted. For the ∼100 Å channel, 12 copies of the AHs, comprising residues S116–L199, were used without inclusion of the TM.

For the ∼52 Å and ∼70 Å channels of *Sc*PEX13, 12 copies of the residues L141–G299, including both the AH and TM regions, were used for the predictions. For the ∼100 Å channel, 12 copies of residues S139–N250, with the TM excluded, were used.

---

**Extended Data Fig. 1.**
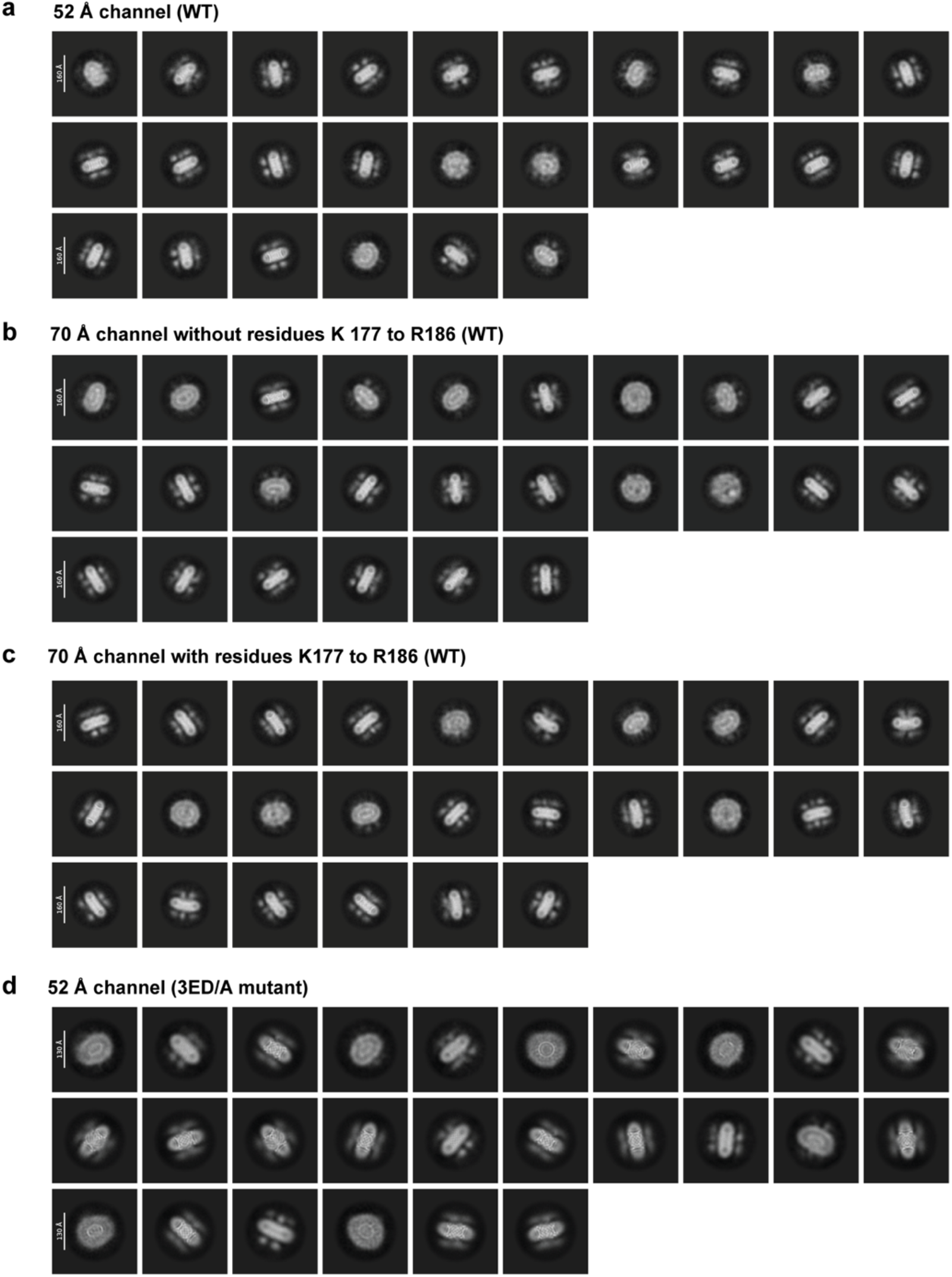
Cryo-EM 2D average classes of different channels. **a**, 2D average classes obtained for the ∼52 Å wild-type (WT) channel. **b**, As in **a**, but for the ∼70 Å wild-type channel lacking density for residues 177-186. **c**, As in **a**, but for the ∼70 Å wild-type channel containing density for residues 177-186. **d**, As in **a**, but for the 3ED/A mutant.

**Extended Data Fig. 2.**
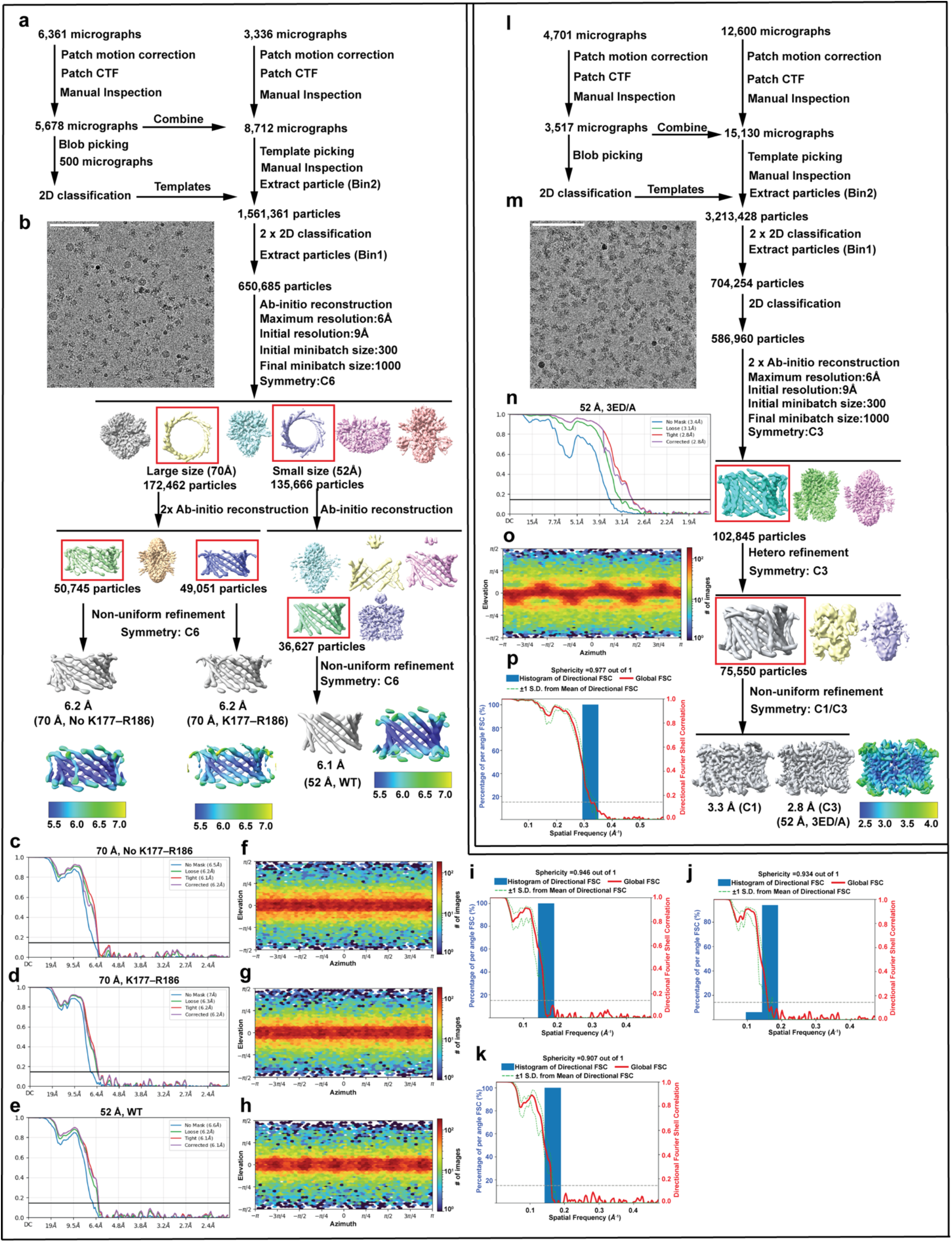
Cryo-EM data processing. **a**, Data-processing scheme for wild-type human PEX13. The particles classes boxed in red were further processed, as indicated. In the final reconstruction, the density map is colored according to local resolution. **b**, Representative raw micrograph. **c**-**e**, Fourier shell correlation (FSC) plots for the ∼70 Å channel map lacking the segment K177–R186, for the ∼70 Å channel map containing the K177–R186 segment, and for the ∼52 Å channel map, respectively. **f**-**h**, Particle orientation distributions for the three wild-type PEX13 maps. **i**-**k**, 3D FSC plots for the three maps. **l**, As in **a**, but for the 3ED/A mutant protein. **m**, As in **b**, but for the 3ED/A mutant protein. **n**, As in **c**-**e**, but for the 3ED/A mutant protein. **o**, As in **f**-**h**, but for the 3ED/A mutant protein. **p**, As in **i**-**k** , but for the 3ED/A mutant protein.

**Extended Data Fig. 3.**
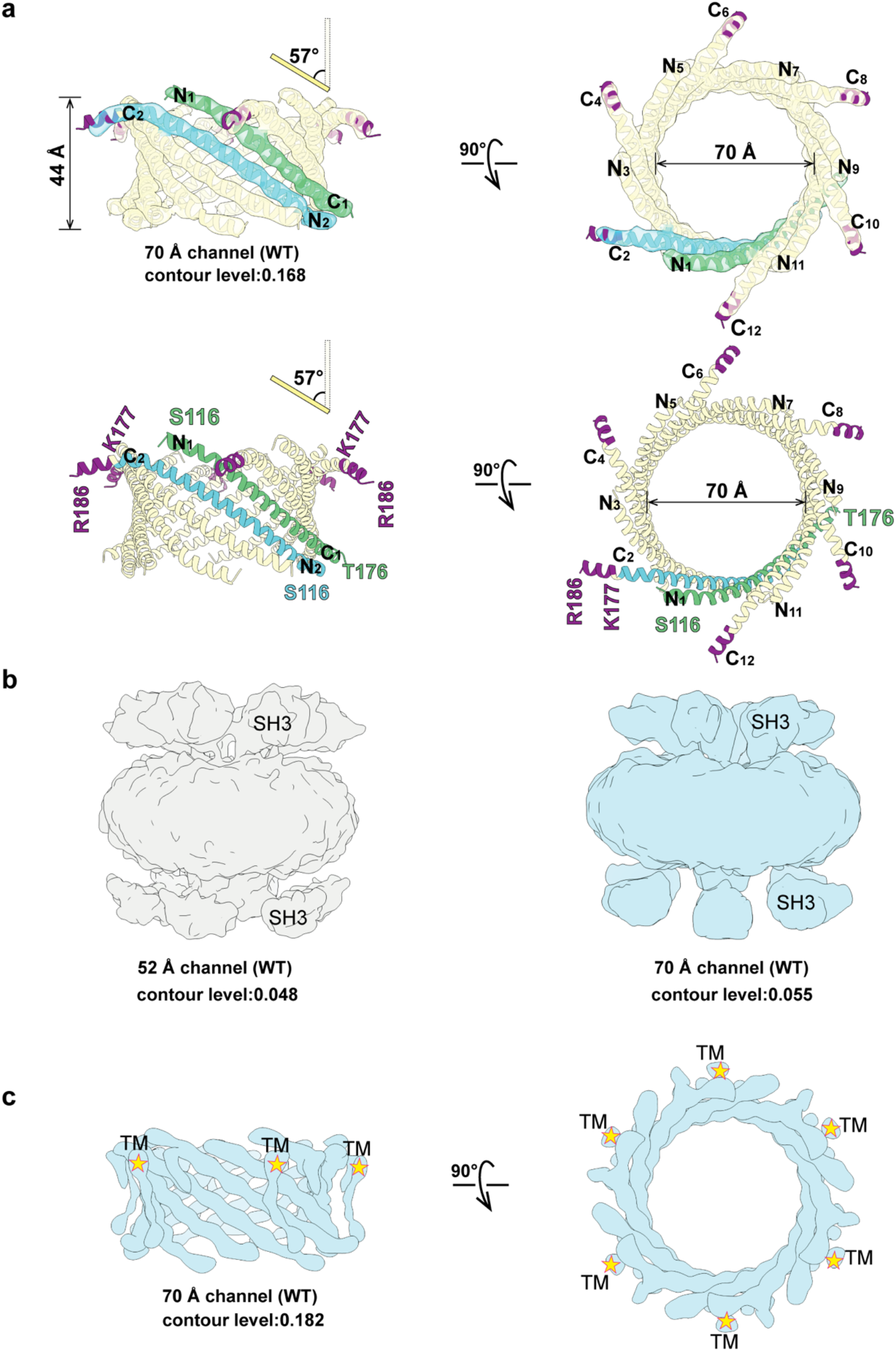
Features in cryo-EM maps of human PEX13. **a**, Semi-transparent density map of the ∼70 Å channel in which residues 177-186 of the AH are visible (shown in purple). Two views are shown in the top panels, with a docked cartoon model based on AlphaFold3 (**Extended Data Fig. 4a**). The lower panels show the model only, with the additional C-terminal residues labeled. **b**, Density maps for the ∼52 Å and ∼70 Å channels at low contour level to visualize the SH3 domains. **c**, Density map for the ∼70 Å channel. Putative density for the TM segments is indicated by a yellow star.

**Extended Data Fig. 4.**
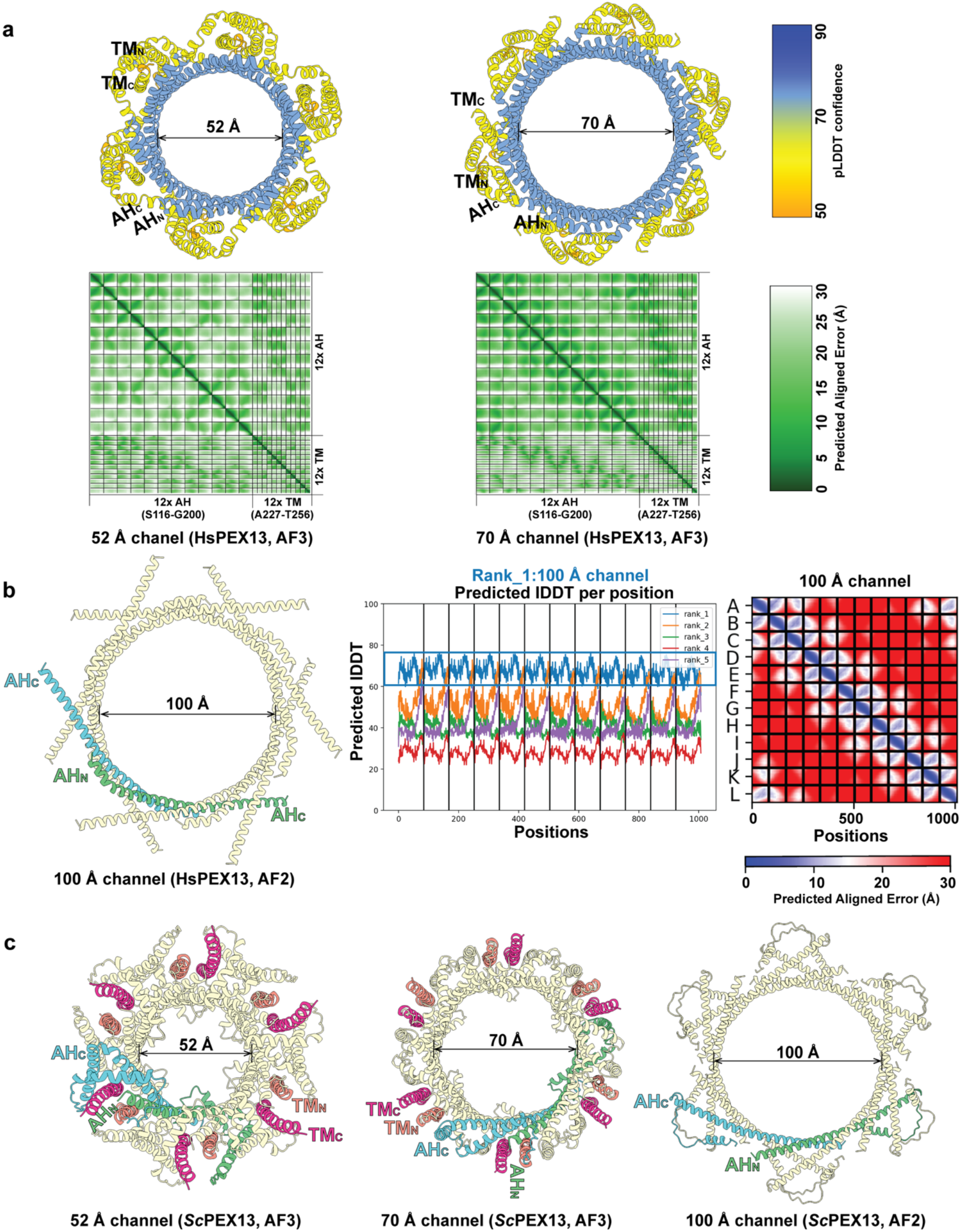
Different channels predicted by AlphaFold. **a**, AlphaFold3 (AF3)-predicted human PEX13 channels with pore diameters of ∼52 Å and ∼70 Å. The predictions were performed with 12 copies of the AH and 12 copies of the TM. The structure is colored according to the pLDDT confidence scale on the right. The lower panels show the predicted aligned error (Å), with the scale on the right. **b**, AlphaFold2 (AF2)-predicted human PEX13 channel with pore diameter of ∼100 Å (left panel), based on rank 1 model that shows the highest values in the pLDDT plot (middle panel). The PAE plot is also shown (right panel), with the color scale at the bottom. **c**, AlphaFold-predicted *S. cerevisiae* PEX13 channels with pore diameters of ∼52 Å, ∼70 Å, and ∼100 Å. The predictions of the smaller channels were done with 12 copies of AH (yellow, green and cyan) and 12 copies of TM (deep pink and salmon). The prediction of the largest channel was done with 12 AH copies only.

**Extended Data Fig. 5.**
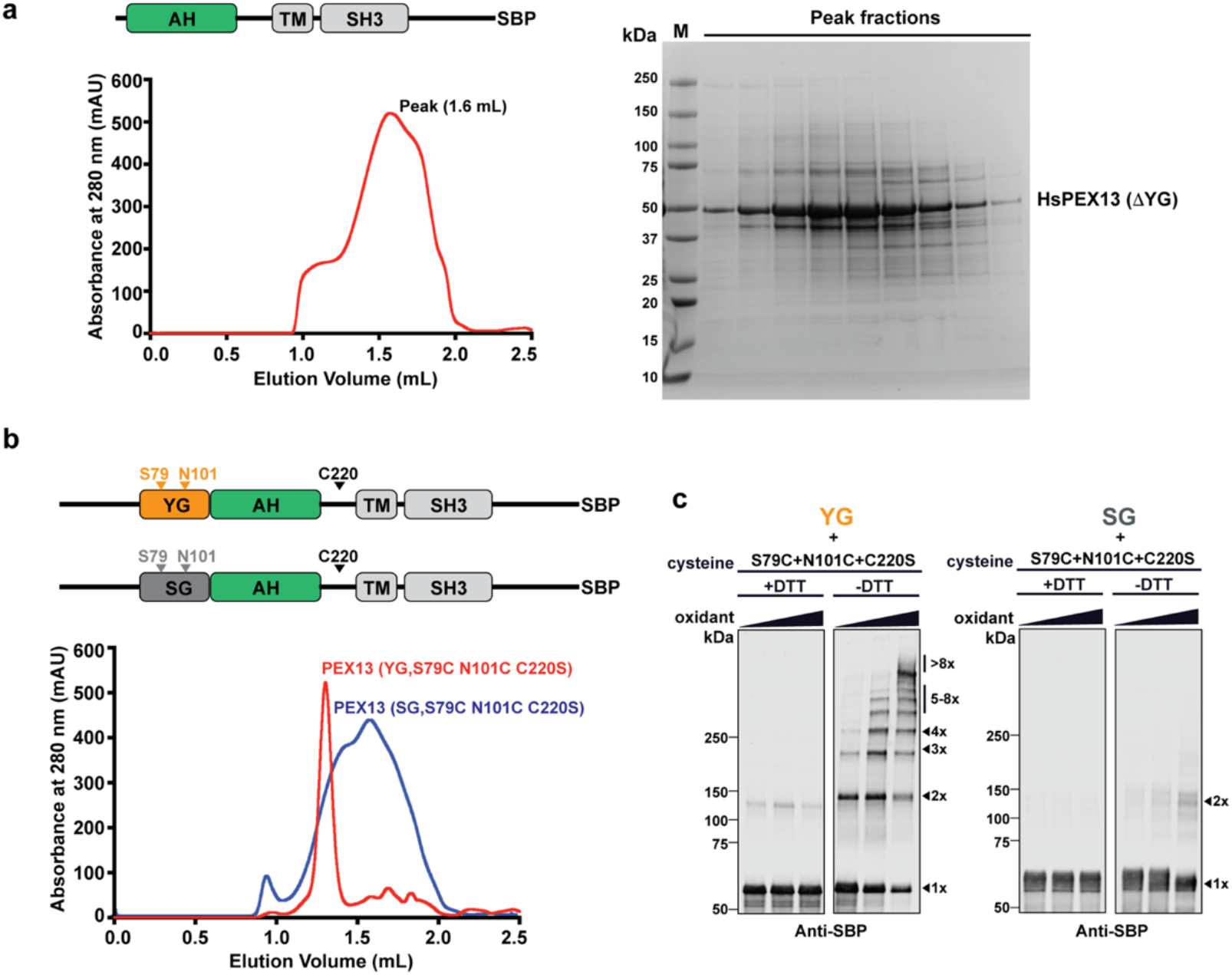
The YG phase is required for the assembly of the PEX13 pore. **a**, SBP-tagged human PEX13 lacking the YG domain (see scheme) was expressed in *P. pastoris*. The protein was solubilized in digitonin, purified with streptavidin beads, and subjected to SEC on a Superose 6 column (left panel). Note that the protein elutes at a lower molecular weight than the wild-type protein (1.6 mL versus 1.3 mL). The peak fractions were analyzed by SDS-PAGE and Coomassie blue staining (right panel). **b**, Human PEX13 with the wild-type YG domain or a mutant in which all tyrosines in the YG domain were mutated to serines (SG mutant) were expressed in *P. pastoris*. Both proteins also contained cysteines incorporated at positions 79 and 101 and had their endogenous cysteine at position 220 changed to serine. The proteins were purified as in (A). Note that the SG mutant (blue curve) elutes in SEC as a broad peak later than the protein with wild-type YG domain (red curve). **c**, The purified proteins were treated with Cu-phenanthroline (oxidant), and analyzed by nonreducing (-DTT) or reducing (+DTT) SDS-PAGE and immunoblotting with SBP antibodies. The number of crosslinked PEX13 molecules in the bands is indicated on the right.

**Extended Data Fig. 6.**
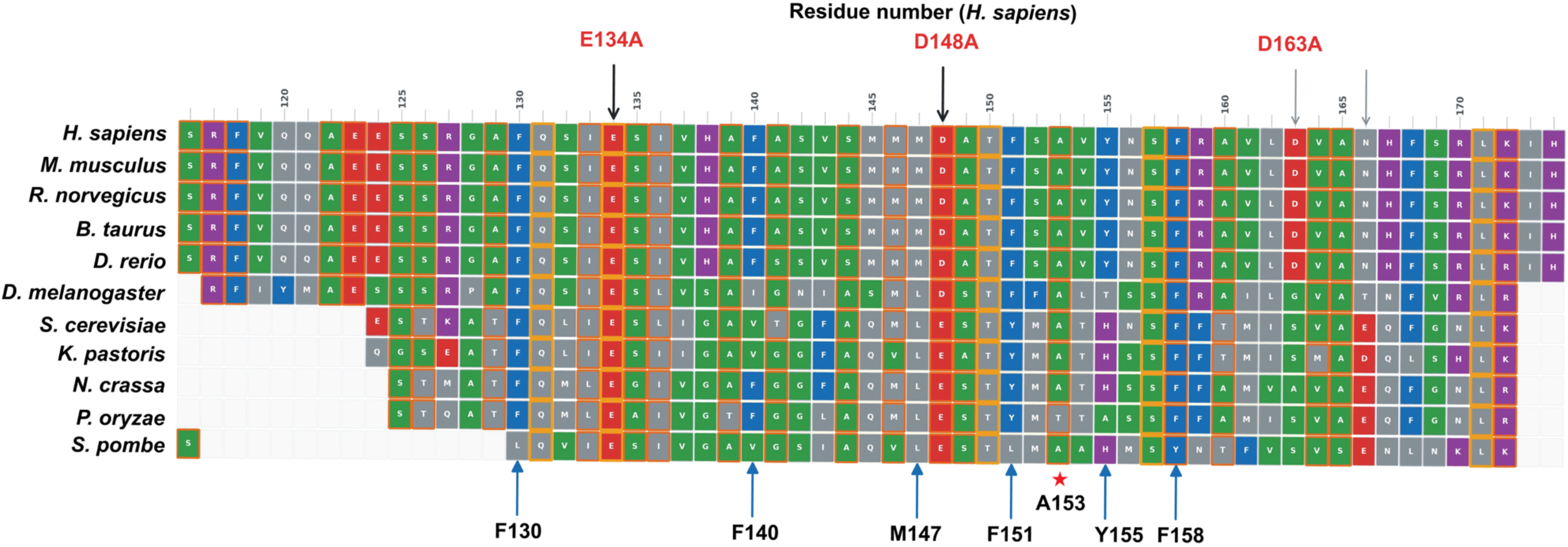
Amino acid sequence conservation of the AH. Sequence alignment of the AHs from organisms representative of different eukaryotic phyla, performed with the help of Claude. The colors group residues by chemical type: red, acidic (D, E); blue, aromatic (F, W, Y, H); purple, basic (K, R); green, small/hydrophilic (A, G, V, S, T); grey, all others. The three highlighted conserved acidic residues were changed to alanines in human PEX13. The corresponding mutations in *S. cerevisiae* PEX13 included the two universally conserved acid residues, as well as a glutamate three residues downstream of D163 that is conserved in some organisms (indicated by an arrow). The blue arrows show the position of amino acids in the constant interface between AHs of human PEX13. The red star indicates the reference residues in the variable interface.

**Extended Data Fig. 7.**
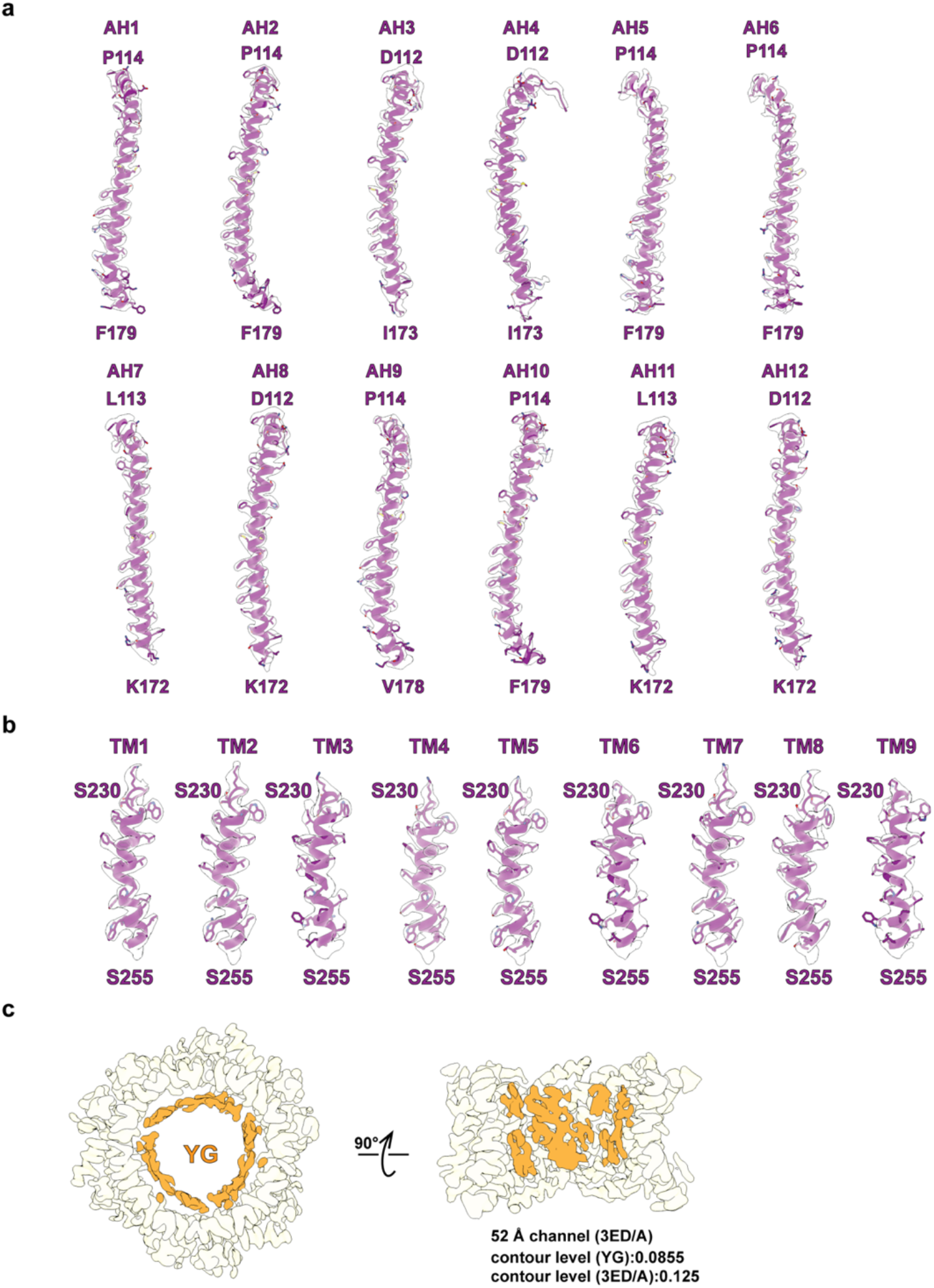
Fit of the model into the density map of the 3ED/A mutant channel. **a**, Shown are the density maps for the 12 AHs and the models built into them. **b**, As in (A), but for the TMs. **c**, Visualization of the YG domains (in orange) at low contour level. The density for the AHs is shown in yellow at a higher threshold level.

**Extended Data Fig. 8.**
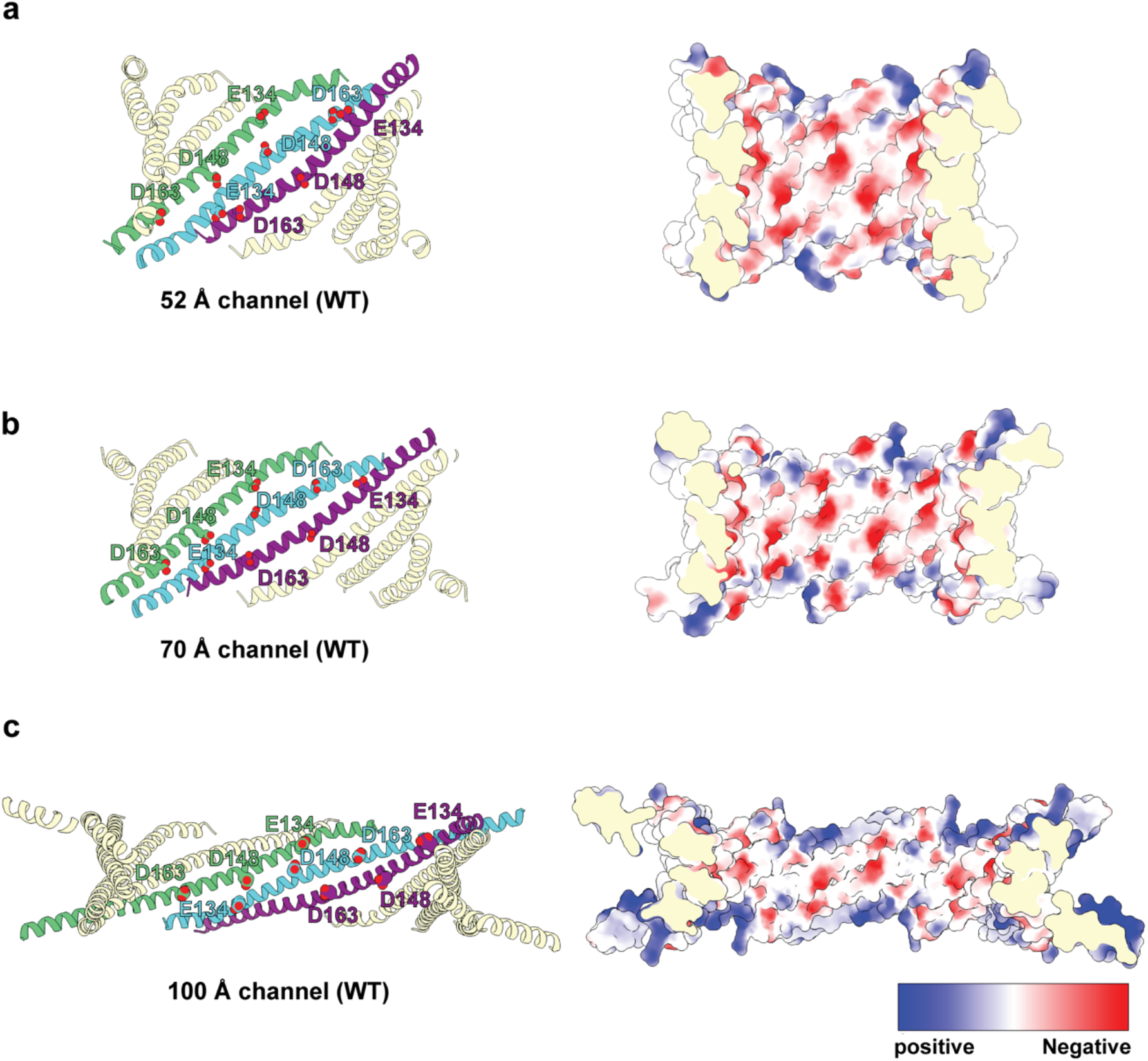
Positions of acidic residues in the human PEX13 channels. **a**, Position of the three conserved acidic residues in the ∼52 Å channel. The left panel shows a cartoon model with the acidic residues marked in three neighboring AHs. The right model shows a space-filling model with the inside of the pore shown in a cut-away view. The surface is colored according to the electrostatic potential of the amino acids (scale at the bottom). **b**, As in **a**, but for the ∼70 Å channel. **c**, As in **a**, but for the ∼100 Å channel. Note that the negative charges are most concentrated in the ∼52 Å channel, which might facilitate the conversion into the active channels by mutual repulsion.

**Extended Data Fig. 9.**
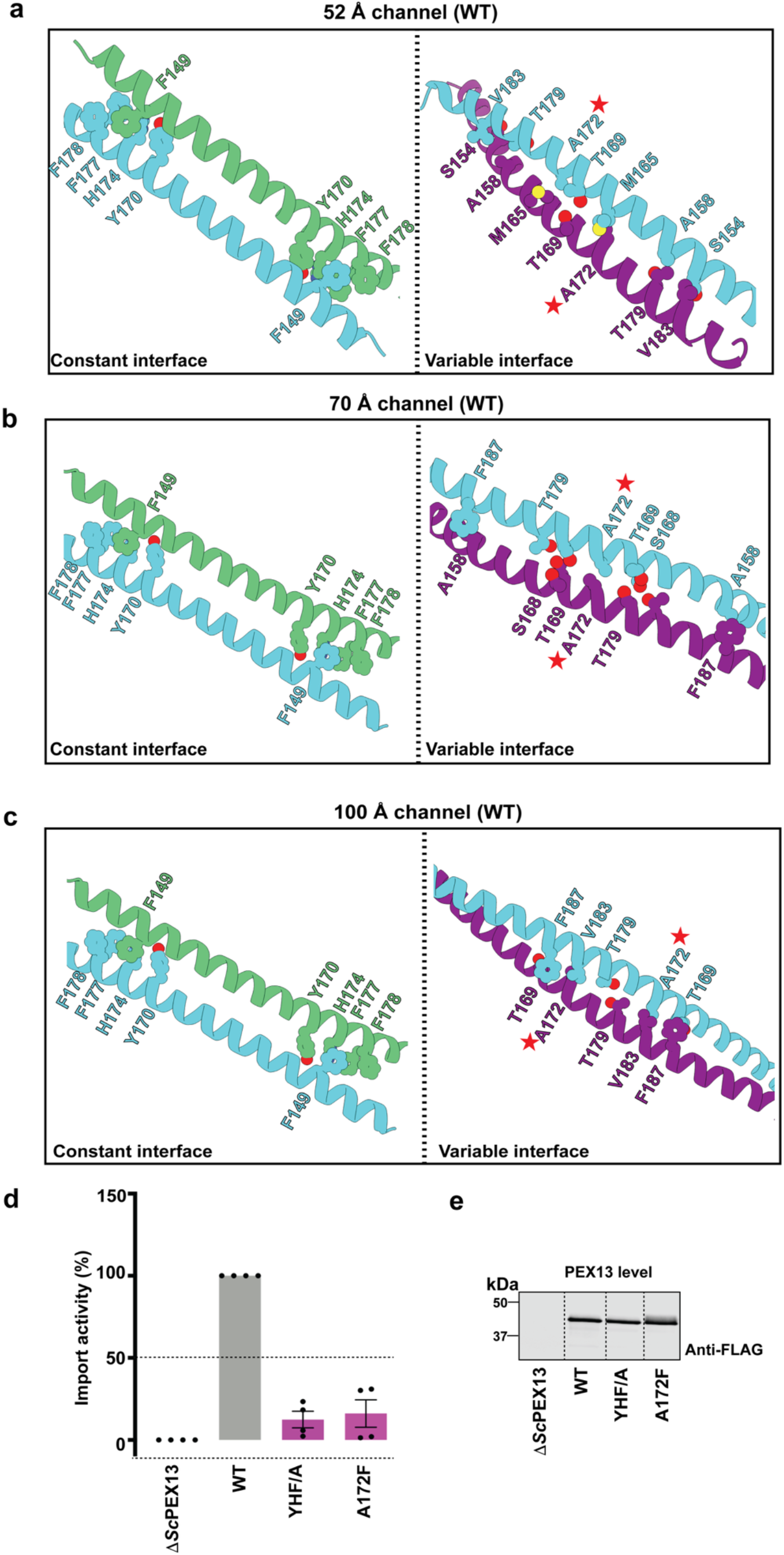
Constant and variable AH interfaces for the predicted *S. cerevisiae* channels. **a**, Constant and variable interfaces of the AlphaFold-predicted ∼52Å channel. Interacting amino acids are labeled and shown with side chains. Hetero-atoms in the side chains are shown in different colors. Note that the constant interface contains aromatic residues, and the variable interface amino acids with small side chains, as in mammals (Fig. 4c). A position in the variable interface is marked by a star to facilitate the comparison with the other channel states shown in **b** and **c**. **b**, As in **a**, but for the predicted ∼70 Å channel. **c**, As in **a**, but for the predicted ∼100 Å channel. **d**. FLAG-tagged wild-type (WT) *S. cerevisiae* PEX13 or the indicated mutants were expressed in *S. cerevisiae* cells lacking endogenous PEX13 (Δ*pex13*). The YHF/A mutant carries alanine mutations of aromatic residues in the constant AH interface (Y170A, H174A, and F177A (corresponding to F151, Y155, and F158 in HsPEX13). The A172F mutant (corresponding to A153 in HsPEX13) carries a mutation in the variable interface. Peroxisomal protein import was measured with an engineered pathway that generates a pigment when import is impaired. Import activity is plotted relative to WT PEX13 (mean ± standard error of 4 experiments). **e**. The expression level of PEX13 in the experiments in **d** was analyzed by SDS-PAGE and immunoblotting with FLAG antibodies.

**Extended Data Fig. 10.**
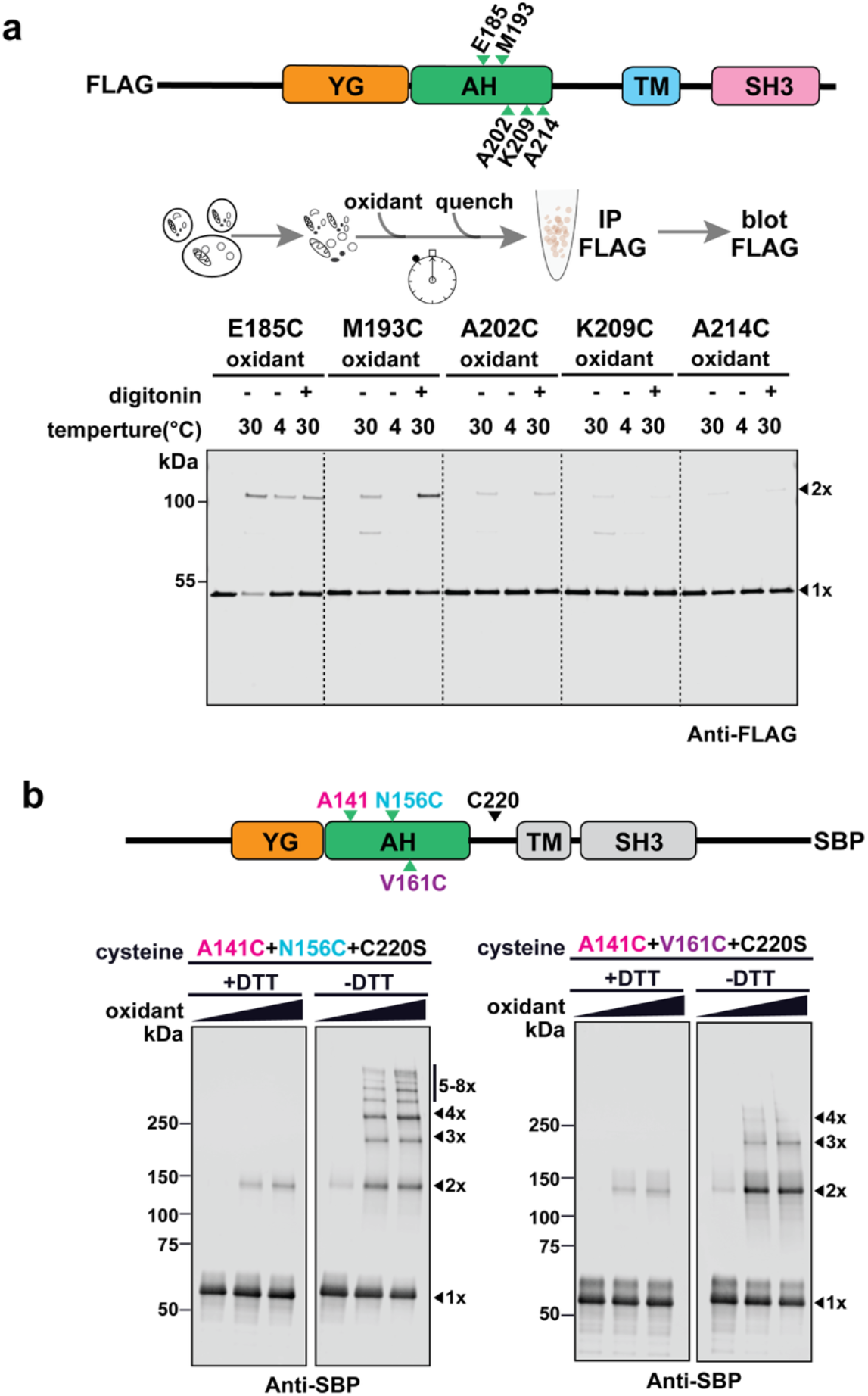
Disulfide crosslinking of yeast and human PEX13. **a**. FLAG-tagged yeast PEX13 variants containing single cysteines at the indicated positions were expressed in *S. cerevisiae* cells. The positions chosen are either in the AH (E185, M193, A202) or in the following region (K209, A214). These positions are not expected to be close to the same position in a neighboring AH and thus serve as controls for the experiment shown in Fig. 4a. **b**, SBP-tagged human PEX13 variants containing two cysteines were expressed in *P. pastoris* cells, purified with streptavidin beads, and treated with an oxidant. The samples were analyzed by reducing (+DTT) or non-reducing (-DTT) SDS-PAGE, followed by immunoblotting with SBP antibodies. One of the cysteines was placed at position A141 in the constant interface of the AHs, the other at position N156 or V161 in the variable interface.

**Extended Data Fig. 11.**
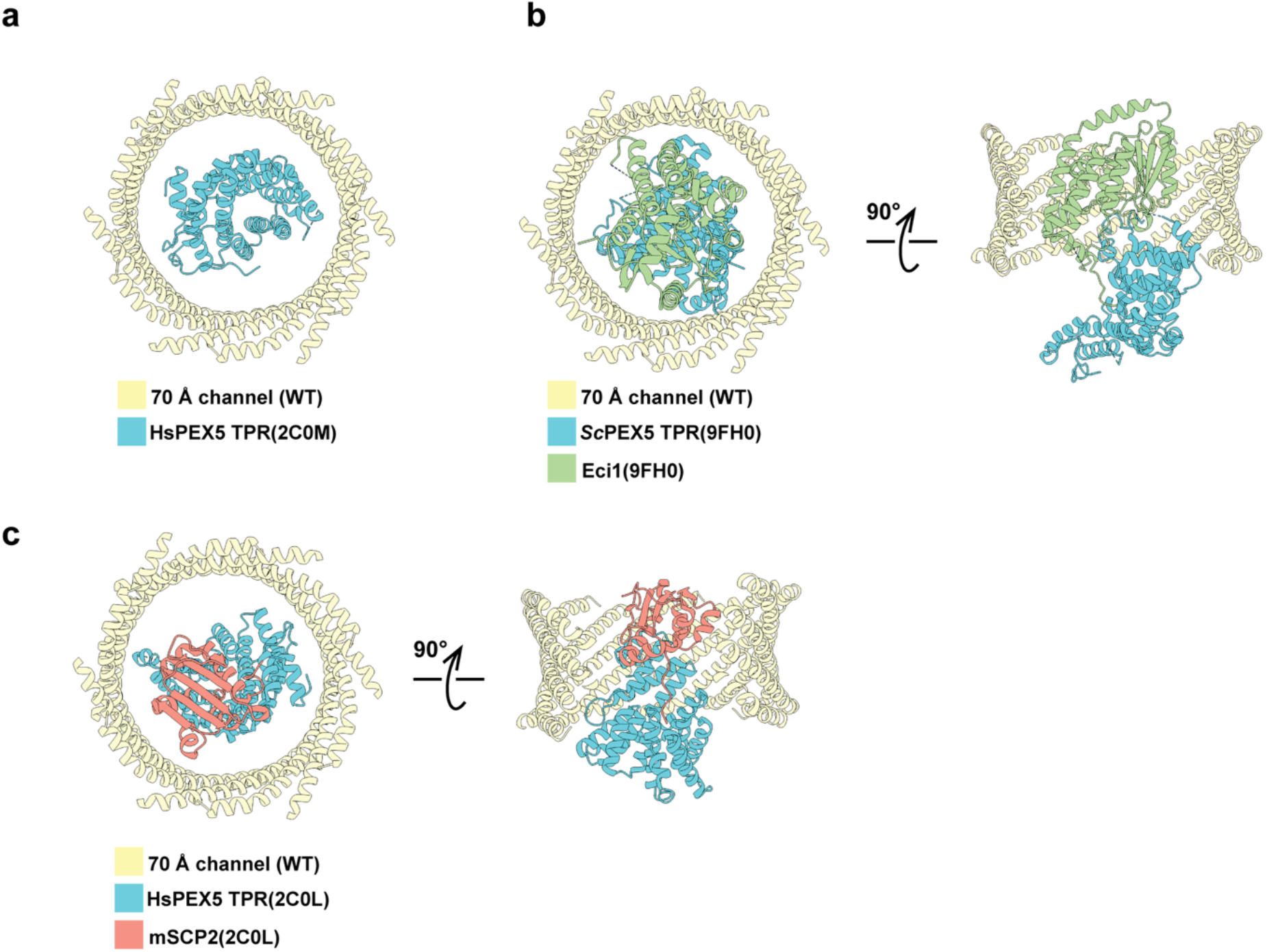
Fitting of receptor and small cargo into the ∼70 Å channel. **a**, The TPR domain of human PEX5 (HsPEX5) (PDB code: 2C0M) was placed into the ∼70 Å channel. Shown is a top view of cartoon models. **b**, As in **a**, but with the complex of the TPR domain of *S. cerevisiae* PEX5 and the cargo protein Eci1 (PDB code: 9FH0). Shown is a top view (left panel) and a side view with the front part of the channel removed for clarity (right panel). **c**, As in **b**, but for the complex of the TPR domain of HsPEX5 and the cargo mSCP2 (PDB code: 2C0L).

**Extended Data Table. 1.** Cryo-EM data collection, refinement and validation statistics in this study.

|  | 52Å import<br>channel (WT)<br>(EMDB:78756) | 70Å import<br>channel (WT, no<br>177-186)<br>(EMB:78762) | 70Å import<br>channel (WT,<br>177-186)<br>(EMB: 78763) | 52Å import<br>channel<br>(3ED/A)<br>(EMDB: 78822)<br>(PDB: 38HQ) |
| --- | --- | --- | --- | --- |
| **Data collection and processing** |  |  |  |  |
| Microscope | Krios | Krios | Krios | Krios |
| Magnification | 81,000 | 81,000 | 81,000 | 81,000 |
| Voltage (kV) | 300 | 300 | 300 | 300 |
| Total electron exposure (e−/Å²) | 50 | 50 | 50 | 50 |
| Defocus range (µm) | -1.0 to -2.5 | -1.0 to -2.5 | -1.0 to -2.5 | -1.0 to -1.8 |
| Pixel size (Å/pixel) | 1.06 | 1.06 | 1.06 | 0.857 |
| Symmetry imposed | C6 | C6 | C6 | C3 |
| Initial particle images (no.) | 1,561,361 | 1,561,361 | 1,561,361 | 3,213,428 |
| Final particle images (no.) | 36,627 | 50,745 | 49,051 | 75,550 |
| Resolution (global, Å) |  |  |  |  |
| FSC 0.5 | 7.2/6.7 | 7.2/6.6 | 7.5/6.8 | 3.9/3.4 |
| (unmasked/masked) | 6.6/6.1 | 6.5/6.2 | 7.0/6.2 | 3.4/2.8 |
| FSC 0.143 |  |  |  |  |
| (unmasked/masked) |  |  |  |  |
| **Model Refinement** |  |  |  |  |
| Initial model used (PDB code) |  |  |  |  |
| Model resolution (Å) |  |  |  | 2.94 |
| FSC threshold |  |  |  | 0.5 |
| Map resolution range (Å) | 5.0 to 8.0 | 5.0 to 8.0 | 5.0 to 8.0 | 2.5 to 5.0 |
| Map sharpening B factor (Å²) | -705.4 | -707.4 | -796.9 | -92.1 |
| Model composition |  |  |  |  |
| Non-hydrogen atoms |  |  |  | 7839 |
| Protein residues |  |  |  | 1003 |
| Ligands |  |  |  | 0 |
| B factors (Å²) |  |  |  |  |
| Protein residues |  |  |  | 79.40 |
| Ligands |  |  |  | --- |
| R.m.s. deviations |  |  |  |  |
| Bond lengths (Å) |  |  |  | 0.002 |
| Bond angles (°) |  |  |  | 0.403 |
| Validation |  |  |  |  |
| MolProbity score |  |  |  | 1.27 |
| Clashscore |  |  |  | 5.03 |
| Poor rotamers (%) |  |  |  | 0.36 |
| Ramachandran plot |  |  |  |  |
| Favored (%) |  |  |  | 99.27 |
| Allowed (%) |  |  |  | 0.73 |
| Disallowed (%) |  |  |  | 0.00 |

**Extended Data Movie 1 Iris-like mechanism of channel conversion**

Shown is the morphing between the PDB files of the ∼52 Å and the ∼70 Å channels (left panels) and between the ∼70 Å and ∼100 Å channels (right panels), generated by Claude. The upper panels show cartoon models of the AHs, viewed down onto the membrane (top views), and the lower panels show side views with AHs in the back removed for clarify. Two dimers connected by a constant interface are shown in orange and blue. Note that the dimers slide relative to one another like an iris, generating the different tilt angles, pore diameters, and pore heights. In the video, all AHs move simultaneously, but in reality the transitions could be more complicated.

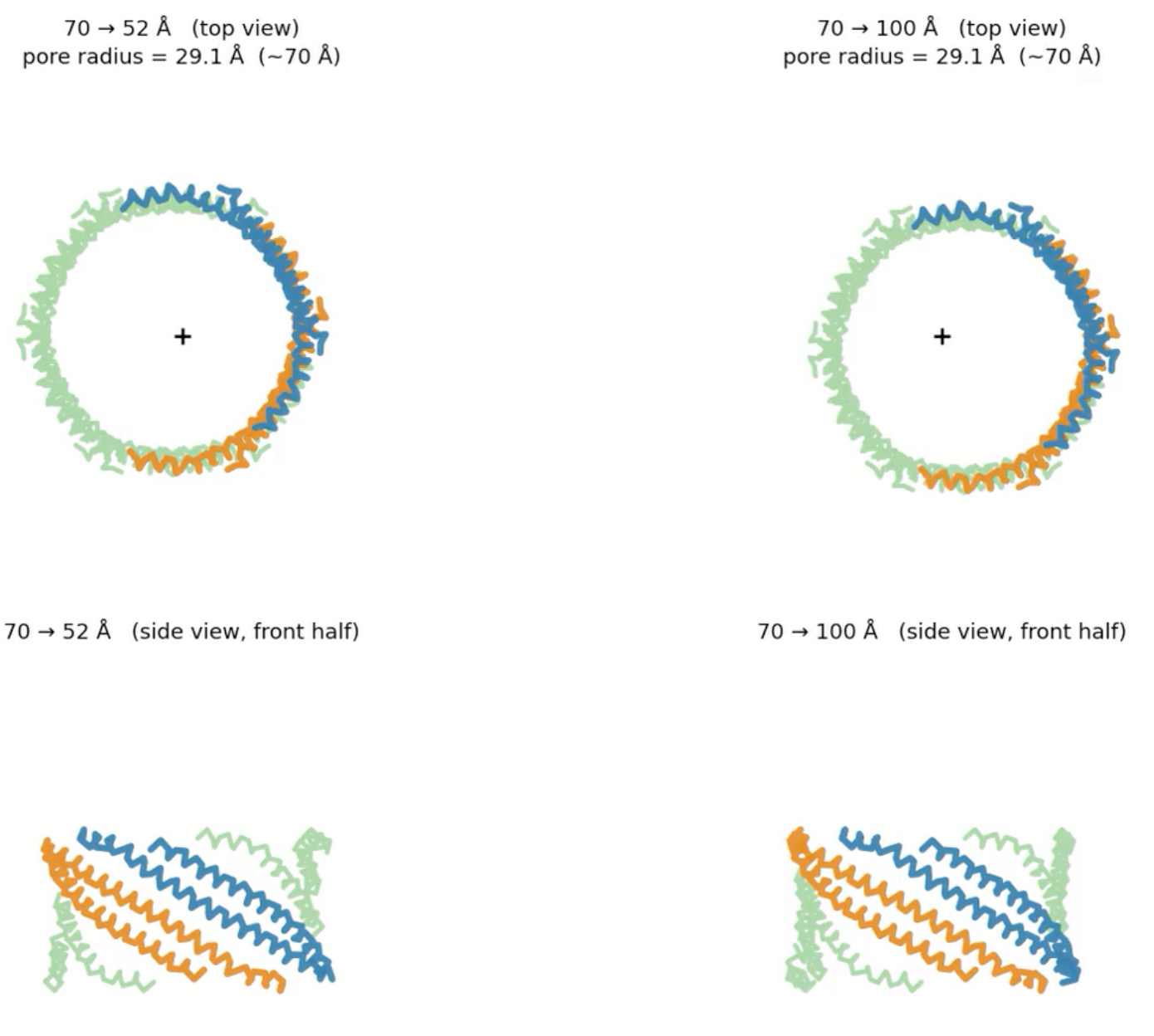

## References

1 Platta, H. W., Jeske, J., Schmidt, N. & Erdmann, R. ATP-Dependent Steps in Peroxisomal Protein Import. Annu Rev Biochem 93, 233–259, doi:10.1146/annurev-biochem-030222-111227 (2024).

2 Francisco, T. et al. Protein transport into peroxisomes: Knowns and unknowns. Bioessays 39, doi:10.1002/bies.201700047 (2017).

3 Fujiki, Y. Peroxisome biogenesis and human peroxisome-deficiency disorders. Proc Jpn Acad Ser B Phys Biol Sci 92, 463–477, doi:10.2183/pjab.92.463 (2016).

4 Jansen, R. L. M., Santana-Molina, C., van den Noort, M., Devos, D. P. & van der Klei, I. J. Comparative Genomics of Peroxisome Biogenesis Proteins: Making Sense of the PEX Proteins. Front Cell Dev Biol 9, 654163, doi:10.3389/fcell.2021.654163 (2021).

5 Farre, J. C., Mahalingam, S. S., Proietto, M. & Subramani, S. Peroxisome biogenesis, membrane contact sites, and quality control. EMBO Rep 20, doi:10.15252/embr.201846864 (2019).

6 Fujiki, Y., Okumoto, K., Mukai, S., Honsho, M. & Tamura, S. Peroxisome biogenesis in mammalian cells. Front Physiol 5, 307, doi:10.3389/fphys.2014.00307 (2014).

7 Walter, T. & Erdmann, R. Current Advances in Protein Import into Peroxisomes. Protein J 38, 351–362, doi:10.1007/s10930-019-09835-6 (2019).

8 Skowyra, M. L., Feng, P. & Rapoport, T. A. Towards solving the mystery of peroxisomal matrix protein import. Trends Cell Biol 34, 388–405, doi:10.1016/j.tcb.2023.08.005 (2024).

9 Purdue, P. E. & Lazarow, P. B. Peroxisome biogenesis. Annu Rev Cell Dev Biol 17, 701–752, doi:10.1146/annurev.cellbio.17.1.701 (2001).

10 Brocard, C. & Hartig, A. Peroxisome targeting signal 1: is it really a simple tripeptide? Biochim Biophys Acta 1763, 1565–1573, doi:10.1016/j.bbamcr.2006.08.022 (2006).

11 Gatto, G. J., Jr., Geisbrecht, B. V., Gould, S. J. & Berg, J. M. Peroxisomal targeting signal-1 recognition by the TPR domains of human PEX5. Nat Struct Biol 7, 1091–1095, doi:10.1038/81930 (2000).

12 Rayapuram, N. & Subramani, S. The importomer--a peroxisomal membrane complex involved in protein translocation into the peroxisome matrix. Biochim Biophys Acta 1763, 1613–1619, doi:10.1016/j.bbamcr.2006.08.035 (2006).

13 Skowyra, M. L. & Rapoport, T. A. PEX5 translocation into and out of peroxisomes drives matrix protein import. Mol Cell 82, 3209–3225 e3207, doi:10.1016/j.molcel.2022.07.004 (2022).

14 Gao, Y., Skowyra, M. L., Feng, P. & Rapoport, T. A. Protein import into peroxisomes occurs through a nuclear pore-like phase. Science 378, eadf3971, doi:10.1126/science.adf3971 (2022).

15 Ravindran, R. et al. Peroxisome biogenesis initiated by protein phase separation. Nature 617, 608–615, doi:10.1038/s41586-023-06044-1 (2023).

16 Schmidt, H. B. & Gorlich, D. Transport Selectivity of Nuclear Pores, Phase Separation, and Membraneless Organelles. Trends Biochem Sci 41, 46–61, doi:10.1016/j.tibs.2015.11.001 (2016).

17 Frey, S., Richter, R. P. & Gorlich, D. FG-rich repeats of nuclear pore proteins form a three-dimensional meshwork with hydrogel-like properties. Science 314, 815–817, doi:10.1126/science.1132516 (2006).

18 Schmidt, H. B. & Gorlich, D. Nup98 FG domains from diverse species spontaneously phase-separate into particles with nuclear pore-like permselectivity. Elife 4, doi:10.7554/eLife.04251 (2015).

19 Feng, P. et al. A peroxisomal ubiquitin ligase complex forms a retrotranslocation channel. Nature 607, 374–380, doi:10.1038/s41586-022-04903-x (2022).

20 Blok, N. B. et al. Unique double-ring structure of the peroxisomal Pex1/Pex6 ATPase complex revealed by cryo-electron microscopy. Proc Natl Acad Sci U S A 112, E4017–4025, doi:10.1073/pnas.1500257112 (2015).

21 Gardner, B. M., Chowdhury, S., Lander, G. C. & Martin, A. The Pex1/Pex6 complex is a heterohexameric AAA+ motor with alternating and highly coordinated subunits. J Mol Biol 427, 1375–1388, doi:10.1016/j.jmb.2015.01.019 (2015).

22 Ciniawsky, S. et al. Molecular snapshots of the Pex1/6 AAA+ complex in action. Nat Commun 6, 7331, doi:10.1038/ncomms8331 (2015).

23 Kunze, M. The type-2 peroxisomal targeting signal. Biochim Biophys Acta Mol Cell Res 1867, 118609, doi:10.1016/j.bbamcr.2019.118609 (2020).

24 Pan, D., Nakatsu, T. & Kato, H. Crystal structure of peroxisomal targeting signal-2 bound to its receptor complex Pex7p-Pex21p. Nat Struct Mol Biol 20, 987–993, doi:10.1038/nsmb.2618 (2013).

25 Skowyra, M. L. & Rapoport, T. A. Import mechanism of peroxisomal proteins with an N-terminal signal sequence. Nat Cell Biol 27, 948–958, doi:10.1038/s41556-025-01662-5 (2025).

26 Chen, W. W. et al. PEX39 facilitates the peroxisomal import of PTS2-containing proteins. Nat Cell Biol 27, 1256–1271, doi:10.1038/s41556-025-01711-z (2025).

27 Stein, K., Schell-Steven, A., Erdmann, R. & Rottensteiner, H. Interactions of Pex7p and Pex18p/Pex21p with the peroxisomal docking machinery: implications for the first steps in PTS2 protein import. Mol Cell Biol 22, 6056–6069, doi:10.1128/MCB.22.17.6056-6069.2002 (2002).

28 Jumper, J. et al. Highly accurate protein structure prediction with AlphaFold. Nature 596, 583–589, doi:10.1038/s41586-021-03819-2 (2021).

29 DeLoache, W. C., Russ, Z. N. & Dueber, J. E. Towards repurposing the yeast peroxisome for compartmentalizing heterologous metabolic pathways. Nat Commun 7, 11152, doi:10.1038/ncomms11152 (2016).

30 Antonenkov, V. D., Sormunen, R. T. & Hiltunen, J. K. The rat liver peroxisomal membrane forms a permeability barrier for cofactors but not for small metabolites in vitro. J Cell Sci 117, 5633–5642, doi:10.1242/jcs.01485 (2004).

31 Stanley, W. A. et al. Recognition of a functional peroxisome type 1 target by the dynamic import receptor pex5p. Mol Cell 24, 653–663, doi:10.1016/j.molcel.2006.10.024 (2006).

32 Peer, L. et al. A cryo-electron microscopy structure of yeast Pex5 in complex with a cargo uncovers a novel binding interface. J Cell Sci 138, doi:10.1242/jcs.263890 (2025).

33 Sridhar, S. et al. Crystallographic binding studies of rat peroxisomal multifunctional enzyme type 1 with 3-ketodecanoyl-CoA: capturing active and inactive states of its hydratase and dehydrogenase catalytic sites. Acta Crystallogr D Struct Biol 76, 1256–1269, doi:10.1107/S2059798320013819 (2020).

34 Evers, M. E., Titorenko, V. I., van der Klei, I. J., Harder, W. & Veenhuis, M. Assembly of alcohol oxidase in peroxisomes of the yeast Hansenula polymorpha requires the cofactor flavin adenine dinucleotide. Mol Biol Cell 5, 829–837, doi:10.1091/mbc.5.8.829 (1994).

35 Waterham, H. R., Russell, K. A., Vries, Y. & Cregg, J. M. Peroxisomal targeting, import, and assembly of alcohol oxidase in Pichia pastoris. J Cell Biol 139, 1419–1431, doi:10.1083/jcb.139.6.1419 (1997).

36 Freitas, M. O. et al. The peroxisomal protein import machinery displays a preference for monomeric substrates. Open Biol 5, 140236, doi:10.1098/rsob.140236 (2015).

37 Fu, L. et al. HIV-1 capsids enter the FG phase of nuclear pores like a transport receptor. Nature 626, 843–851, doi:10.1038/s41586-023-06966-w (2024).

38 Kreysing, J. P. et al. Passage of the HIV capsid cracks the nuclear pore. Cell 188, 930–943 e921, doi:10.1016/j.cell.2024.12.008 (2025).

39 Lee, M. S., Mullen, R. T. & Trelease, R. N. Oilseed isocitrate lyases lacking their essential type 1 peroxisomal targeting signal are piggybacked to glyoxysomes. Plant Cell 9, 185–197, doi:10.1105/tpc.9.2.185 (1997).

40 Walton, P. A., Hill, P. E. & Subramani, S. Import of stably folded proteins into peroxisomes. Mol Biol Cell 6, 675–683, doi:10.1091/mbc.6.6.675 (1995).

41 Gaussmann, S. et al. Modulation of peroxisomal import by the PEX13 SH3 domain and a proximal FxxxF binding motif. Nat Commun 15, 3317, doi:10.1038/s41467-024-47605-w (2024).

42 Williams, C. & Distel, B. Pex13p: docking or cargo handling protein? Biochim Biophys Acta 1763, 1585–1591, doi:10.1016/j.bbamcr.2006.09.007 (2006).

43 Douangamath, A. et al. Topography for independent binding of alpha-helical and PPII-helical ligands to a peroxisomal SH3 domain. Mol Cell 10, 1007–1017, doi:10.1016/s1097-2765(02)00749-9 (2002).

44 Rapoport, T. A., Li, L. & Park, E. Structural and Mechanistic Insights into Protein Translocation. Annu Rev Cell Dev Biol 33, 369–390, doi:10.1146/annurev-cellbio-100616-060439 (2017).

45 Araiso, Y., Imai, K. & Endo, T. Role of the TOM Complex in Protein Import into Mitochondria: Structural Views. Annu Rev Biochem 91, 679–703, doi:10.1146/annurev-biochem-032620-104527 (2022).

46 Liu, H., Li, A., Rochaix, J. D. & Liu, Z. Architecture of chloroplast TOC-TIC translocon supercomplex. Nature 615, 349–357, doi:10.1038/s41586-023-05744-y (2023).

47 Chen, Y. et al. YidC Insertase of Escherichia coli: Water Accessibility and Membrane Shaping. Structure 25, 1403–1414 e1403, doi:10.1016/j.str.2017.07.008 (2017).

48 Kumazaki, K. et al. Structural basis of Sec-independent membrane protein insertion by YidC. Nature 509, 516–520, doi:10.1038/nature13167 (2014).

49 Wu, X. et al. Structural basis of ER-associated protein degradation mediated by the Hrd1 ubiquitin ligase complex. Science 368, doi:10.1126/science.aaz2449 (2020).

50 Wu, X. & Rapoport, T. A. Translocation of Proteins through a Distorted Lipid Bilayer. Trends Cell Biol 31, 473–484, doi:10.1016/j.tcb.2021.01.002 (2021).

51 Leon, S., Goodman, J. M. & Subramani, S. Uniqueness of the mechanism of protein import into the peroxisome matrix: transport of folded, co-factor-bound and oligomeric proteins by shuttling receptors. Biochim Biophys Acta 1763, 1552–1564, doi:10.1016/j.bbamcr.2006.08.037 (2006).

52 Mueller, M., Grauschopf, U., Maier, T., Glockshuber, R. & Ban, N. The structure of a cytolytic alpha-helical toxin pore reveals its assembly mechanism. Nature 459, 726–730, doi:10.1038/nature08026 (2009).

53 Tanaka, K., Caaveiro, J. M., Morante, K., Gonzalez-Manas, J. M. & Tsumoto, K. Structural basis for self-assembly of a cytolytic pore lined by protein and lipid. Nat Commun 6, 6337, doi:10.1038/ncomms7337 (2015).

54 Zimmerli, C. E. et al. Nuclear pores dilate and constrict in cellulo. Science 374, eabd9776, doi:10.1126/science.abd9776 (2021).

## References

55 Gietz, R. D. & Schiestl, R. H. High-efficiency yeast transformation using the LiAc/SS carrier DNA/PEG method. Nat Protoc 2, 31–34, doi:10.1038/nprot.2007.13 (2007).

56 Punjani, A., Rubinstein, J. L., Fleet, D. J. & Brubaker, M. A. cryoSPARC: algorithms for rapid unsupervised cryo-EM structure determination. Nat Methods 14, 290–296, doi:10.1038/nmeth.4169 (2017).

57 Punjani, A., Zhang, H. & Fleet, D. J. Non-uniform refinement: adaptive regularization improves single-particle cryo-EM reconstruction. Nat Methods 17, 1214–1221, doi:10.1038/s41592-020-00990-8 (2020).

58 Tan, Y. Z. et al. Addressing preferred specimen orientation in single-particle cryo-EM through tilting. Nat Methods 14, 793–796, doi:10.1038/nmeth.4347 (2017).

59 Morin, A. et al. Collaboration gets the most out of software. Elife 2, e01456, doi:10.7554/eLife.01456 (2013).

60 Pettersen, E. F. et al. UCSF ChimeraX: Structure visualization for researchers, educators, and developers. Protein Sci 30, 70–82, doi:10.1002/pro.3943 (2021).

61 Emsley, P. & Cowtan, K. Coot: model-building tools for molecular graphics. Acta Crystallogr D Biol Crystallogr 60, 2126–2132, doi:10.1107/S0907444904019158 (2004).

62 Afonine, P. V. et al. Real-space refinement in PHENIX for cryo-EM and crystallography. Acta Crystallogr D Struct Biol 74, 531–544, doi:10.1107/S2059798318006551 (2018).

